# Transcriptional modulation unique to vulnerable motor neurons predicts ALS across species and SOD1 mutations

**DOI:** 10.1101/2024.03.15.584775

**Authors:** Irene Mei, Susanne Nichterwitz, Melanie Leboeuf, Jik Nijssen, Isadora Lenoel, Dirk Repsilber, Christian S. Lobsiger, Eva Hedlund

## Abstract

Amyotrophic lateral sclerosis (ALS) is characterized by the progressive loss of motor neurons that innervate skeletal muscles. However, certain motor neuron groups including ocular motor neurons, are relatively resilient. To reveal key drivers of resilience versus vulnerability in ALS, we investigate the transcriptional dynamics of four distinct motor neuron populations in SOD1G93A ALS mice using LCM-seq and single molecule fluorescent *in situ* hybridization. We find that resilient ocular motor neurons regulate few genes in response to disease. Instead, they exhibit high baseline gene expression of neuroprotective factors including *En1*, *Pvalb, Cd63* and *Gal,* some of which vulnerable motor neurons upregulate during disease. Vulnerable motor neuron groups upregulate both detrimental and regenerative responses to ALS and share pathway activation, indicating that breakdown occurs through similar mechanisms across vulnerable neurons, albeit with distinct timing. Meta-analysis across four rodent mutant SOD1 motor neuron transcriptome datasets identify a shared vulnerability code of 39 genes including e.g *Atf4*, *Nupr1*, *Ddit3 and Penk,* involved in apoptosis, as well as a proregenerative and anti-apoptotic signature consisting of *Atf3*, *Vgf*, *Ina*, *Sprr1a, Fgf21, Gap43, Adcyap1,* and *Mt1*. Machine learning using genes upregulated in SOD1G93A spinal motor neuron predicts disease in human stem cell-derived *SOD1E100G* motor neurons, and shows that dysregulation of *VGF*, *INA, PENK* and *NTS* are strong disease-predictors across species and SOD1 mutations. Our study reveals motor neuron population-specific gene expression and temporal disease-induced regulation that together provide a basis to explain ALS selective vulnerability and resilience and that can be used to predict disease.

## Introduction

Amyotrophic lateral sclerosis (ALS) is a neurodegenerative disease that is characterized by the loss of upper motor neurons in the cortex, leading to spasticity, and lower somatic motor neurons in the brainstem and spinal cord that control skeletal muscles, leading to muscle atrophy and paralysis. About 10% of cases are familial and of these, around 20% are caused by dominant mutations in the superoxide dismutase 1 (SOD1) gene, which induce ALS through gain-of-toxic function mechanisms (Rosen et al. 1993). SOD1 is ubiquitously expressed in the body and when mutated it accumulates across cell types over time. Mutant SOD1 affects the function of many cell types and yet it causes selective loss of particular neurons only. In fact, not even all somatic motor neurons are affected in ALS, but certain subpopulations show a strong resilience, including the oculomotor (cranial nerve 3, CN3), trochlear (CN4) and abducens (CN6) motor neurons, which innervate the extraocular muscles that control eye movement. This has been demonstrated both in sporadic ALS patients (Caligari et al. 2013; Gizzi et al. 1992; Kubota et al. 2000), in mutant SOD1 ALS mice (Comley et al. 2016; Tjust et al. 2012; Valdez et al. 2013) and in inducible mutant TDP-43 mice (Spiller et al. 2016). Thus, this resilience is present across disease causations and species. Furthermore, visceral motor neurons that innervate smooth muscle appear relatively unaffected in ALS patients (Shimizu et al. 2011; McLeod et al. 2022). Why particular neurons are more resilient than others to ALS and if this is entirely intrinsically programmed or also due to differences in non-cell autonomous factors in the environments surrounding motor neurons or in their muscle targets is still unclear. Through removing mutant SOD1 from particular cell types in transgenic mice, it has been demonstrated that mutant SOD1 within motor neurons themselves is important for driving initiation and early progression of disease (Boillee et al. 2006), while microglia and astrocytes are important for later disease progression stages (Boillee et al. 2006; Yamanaka et al. 2008). Stem cell-derived oculomotor neurons also show a higher level of resistance to ALS-like toxicity *in vitro* indicating that part of their resilience comes from within (Allodi et al. 2018). Furthermore, we have demonstrated that oculomotor neurons express relatively high levels of different growth factors at baseline and in response to disease, as shown for spinal muscular atrophy (SMA), which can protect motor neurons and their synapses with muscle (Allodi et al. 2016; Nichterwitz et al. 2020). Oculomotor neurons also lack baseline expression of Mmp9, which particularly vulnerable spinal motor neurons express high levels of and which appears detrimental to motor neurons (Kaplan et al. 2013). Thus, it is likely the presence as well as lack of certain factors within neurons at baseline, in combination with their response to disease, that render a cell particularly resilient versus vulnerable to disease-stressors. Several studies have analyzed the transcriptional dysregulation that occurs within vulnerable spinal motor neurons in mutant SOD1 mouse models of ALS either prior to overt clinical symptoms (Perrin et al. 2005; Lobsiger et al. 2007), or at symptomatic stages (Perrin et al. 2005; Sun et al. 2015; Shadrach et al. 2021). These studies have shown that vulnerable neurons respond to disease by activating pathways of compensatory regeneration and response to DNA damage and cell injury.

No attempts have yet been made to comprehensively analyze how different types of vulnerable neurons respond to ALS over time in comparison to different types of relatively resilient neurons, in order to disseminate the mechanisms by which particular neurons are vulnerable while others remain relatively resilient. We reasoned that an analysis of how both vulnerable and resilient neurons respond to mutant SOD1 over time, as well as careful consideration of their baseline gene expression, could give further insight into disease mechanisms. We therefore set out to analyze different types of resilient motor neurons, including somatic oculomotor (CN3) and trochlear (CN4) motor neurons, and visceral motor neurons of the dorsal motor vagus nerve (CN10) to delineate if all resilient neurons respond in a similar fashion to ALS. We compared two vulnerable motor neuron populations; lumbar spinal motor neurons and hypoglossal motor neurons (CN12), to also answer the question if vulnerable neurons share destructive pathways and attempts of compensatory responses or if the brainstem and spinal cord are going through distinct disease mechanisms, due to differences in target innervation and local environments. We used laser capture microdissection and Smart-seq2 RNA sequencing (LCM-seq) (Nichterwitz et al. 2016, 2018) to analyze the transcriptional dynamics across neuronal populations in ALS mice and wildtype littermates and confirmed differential gene expression (DEGs) using RNAscope. We also conducted meta-analysis on existing transcriptomic datasets on rodent mutant SOD1 spinal motor neurons and machine learning and to identify strong disease predictors and a general vulnerability code across mutations and species.

## Results

### Transcriptional regulation of mutant SOD1 or other disease associated genes do not underlie selective vulnerability among motor neurons

To retrieve temporal mechanistic insight into the selective vulnerability and resilience across motor neuron populations in ALS we isolated somatic motor neurons from CN3/4 (resilient), CN12 (vulnerable), and lumbar spinal cord (highly vulnerable) as well as visceral motor neurons from CN10 (resilient) of presymptomatic (P56) and onset of symptoms (P112) SOD1G93A mice and wild-type littermates using laser capture microdissection. We subjected the neurons to Smart-Seq2 polyA-based RNA sequencing (Fig. 1A, Supplemental Fig. S1A-D’’). Reads uniquely mapped to mouse genome (69.7 ± 0.50% mean±SEM) were used for further analyses (Supplemental Fig. S2A). Pairwise Pearson correlation of the variance stabilized transformed (VST) expression values of all samples that passed the quality control (see *Methods)* shows a Pearson correlation of at least 0.95 within each cell type and did not show any separation of disease versus control within the respective cell types (Fig. 1B).

**Fig. 1.**
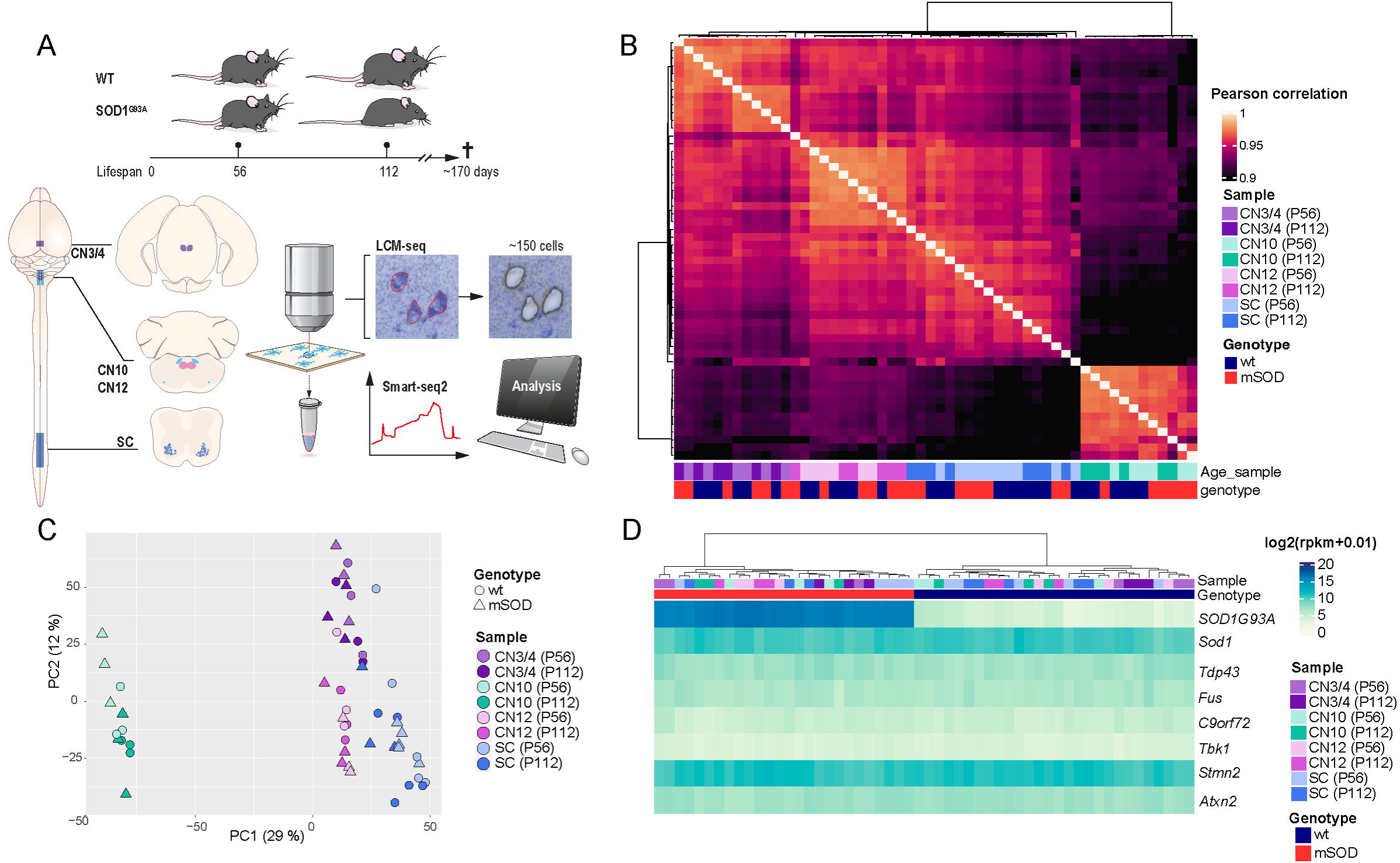
Motor neuron subpopulations show unique transcriptional profiles but similar levels of ALS disease gene expression. **A**. Schematic representation of the study design and LCM-seq workflow. We used SOD1G93A mice, a well-established ALS model, and their wild-type (WT) littermates as controls at postnatal day 56 (P56, presymptomatic) and postnatal day 112 (P112, disease onset). Laser capture microdissection (LCM) was performed to isolate motor neurons from three cranial nerve (CN) nuclei (CN3/4, CN10, CN12) and spinal cord (SC), followed by Smart-seq2 RNA sequencing for transcriptome analysis. **B**. Pairwise Pearson’s correlation heatmap of variance-stabilized transformed (VST) gene expression data, showing hierarchical clustering of samples by cell type, genotype, and age. Single linkage method was used for the hierarchical clustering of columns. **C**. Principal component analysis (PCA) based on whole transcriptome expression data highlights clear separation between different motor neuron subtypes, with genotype and age also influencing clustering. **D**. of key ALS-associated genes (*Sod1*, *Tardp43*, *Fus*, *C9orf72*, *Tbk1*, *Stmn2*, *Atxn2*), showing no major differences across motor neuron populations. Abbreviations: (P) Postnatal day; (LCM-seq) laser capture microdissection coupled with RNA sequencing; (SC) spinal cord; (CN) cranial nerve; (CN12) hypoglossal nucleus; (CN10) dorsal motor nucleus of the vagus nerve; (CN3/4) oculomotor and trochlear nuclei.

Hierarchical clustering of samples based on *Phox2a/b* and *Hox* gene expression shows a clustering based on cell type and position along the anterior-posterior body axis and confirms that CN3/4 neurons cluster based on *Phox2b* expression and lack of *Hox* gene expression, while spinal motor neurons clustered based on caudal *Hox* gene expression (*Hox6-11*) compared to CN10 and CN12 neurons, as expected (Supplemental Fig. S2B). Further marker gene expression analysis showed that all isolated motor neuron groups expressed *Chat* and *Islet1* (*Isl1*). CN12 and spinal motor neurons also expressed *Isl2* and *Mnx1*, while those markers were nearly absent in CN3/4 and visceral CN10 motor neurons (Supplemental Fig. S2C). All neuron groups expressed high levels of peripherin (*Prph*) and neurofilament heavy chain (*Nefh*), although CN10 had markedly lower levels of *Nefh* than the other groups (Supplemental Fig. S2C). Principal component analysis including all expressed genes showed that samples cluster based on cell type (Fig. 1C). The first principal component (PC1) clearly distinguished visceral (CN10) from the somatic (CN3/4, CN12 and spinal) motor neurons. The PC1 also to a lesser extent distinguished brainstem motor neurons from spinal motor neurons, while PC2 distinguished CN3/4 motor neurons from the other groups (Fig. 1C). Within the CN10 group, the PC2 axis also seemed to resolve age, but this was not the case for any other neuron group. ALS neurons were not clearly separated from control neurons in this plot of all expressed genes (Fig. 1C), as a minority of genes were dysregulated in ALS compared to the overall gene expression across these neurons (Fig. 2A). Analysis of additional PCs (PC3-6) did not further resolve diseases versus control (Supplemental Fig. S2F-G). The ALS mouse model is based on overexpression of human mutated *SOD1G93A* and the level of the resulting protein is important for disease development. Analysis of human and mouse *Sod1* levels in mutant SOD1G93A mice and wildtype littermates clearly shows that human *SOD1G93A* mRNA is highly expressed in the SOD1G93A mice across neuron types and that the endogenous mouse *Sod1* mRNA is expressed, albeit at lower level than mutant*SOD1G93A*, across all mice, wild-type and SOD1G93A, as expected (Fig. 1D). As the level of mutant *SOD1* is an important disease predictor we conducted a careful statistical analysis of any variation in expression level across cell types. We find only very minor variations in *SOD1G93A* levels across cell types, with CN10 and CN12 motor neurons showing the highest levels. Furthermore, mutant *SOD1 mRNA* levels did not change significantly with age and thus do not explain onset of disease across the neuron types (Supplemental Fig. S2F). We conducted an ANOVA analysis comparing the two homologs of the *Sod1* gene, followed by a post-hoc Bonferroni-corrected t-test. The pairwise comparisons revealed significant but subtle differences between groups (Supplemental Fig.S2F). Collectively, these small differences in mRNA level of *SOD1G93A* do not reflect the level of vulnerability of the respective cell types. Analysis of endogenous mouse *Sod1* mRNA confirmed that it was expressed at a lower level than human *SOD1G93A*, and demonstrated that there was only a very slight difference in levels across cell types (Supplemental Fig. S2F). Thus, differential regulation of transcripts other than *SOD1G93A* and *Sod1* hold the key to understanding differential vulnerability in SOD-ALS. Analysis of mouse embryonic stem cell derived neurons has indicated that cranial motor neurons have higher proteasome activity to degrade misfolded proteins compared to spinal motor neurons and thus would be more resilient to proteostatic stress and misfolded SOD1 (An et al. 2019). To investigate this matter *in vivo*, we analyzed GO terms related to proteasome, ubiquitination and proteolysis across samples of CN3/4 and spinal motor neurons in control and SOD1G93A mice, using Gene Set Variation Analysis (Hänzelmann S. et al. 2013, GSVA). We found no indication on the transcriptome level that CN3/4 motor neurons would have a better capacity to degrade proteins than spinal motor neurons in either control or SOD1G93A mice (Supplemental Fig. S2G), but this does not exclude translational or posttranslational regulation of proteasome activity. We subsequently analyzed the expression levels of other disease-causing or disease-modifying genes, including *Tardbp*, *Fus*, *C9orf72*, *Tbk1*, *Stmn2* and *Atxn2*, but found no variation across either cell types or disease states across cell types (Fig. 1D). In conclusion, different motor neuron types are clearly distinguished by their transcriptomic profiles and *SOD1G93A*-induced gene expression changes do not shift the identity of the motor neurons in any major way. *SOD1G93A* expression levels were comparable across neuron types, and thus do not reflect differences in susceptibility, neither does the expression level of other disease-causative or modifying genes. Therefore, to understand selective vulnerability, further analysis of genes and pathways dysregulated downstream of mutant *SOD1G93A* may reveal how cells respond differently and thus either succumb to or are shielded from disease.

**Fig. 2.**
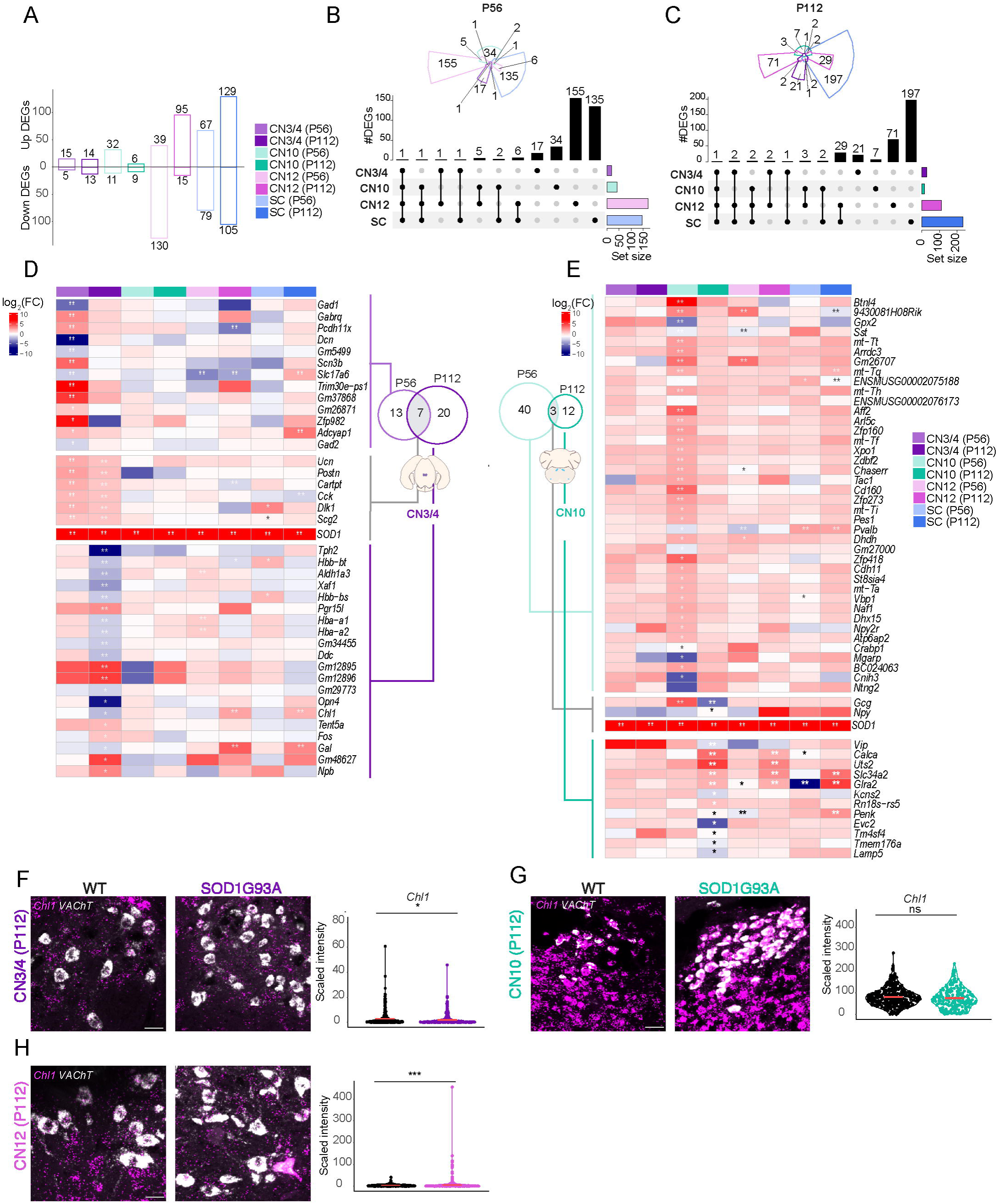
Relatively resistant motor neurons show little gene dysregulation in response to mutant SOD1. (**A**). Bar plots of differentially expressed genes (DEGs) across motor neuron populations in SOD1G93A versus WT mice, highlighting a low number of DEGs in CN3/4 compared to other populations. (**B-C**) Upset plots of DEGs at P56 and P112, showing the number of genes differentially expressed across CN3/4, CN10, CN12, and SC at different disease stages (Likelihood ratio test, Benjamini-Hochberg adjusted p-value < 0.05). (**D**) Heatmap showing log fold change (logFC) values of the DEGs across disease stages for CN3/4 at both P56 and P112, showing relatively few significant changes compared to other populations. (Likelihood ratio test, ** Benjamini Hochberg FDR < 0.01; * Benjamini Hochberg FDR < 0.05). (**E)** Heatmaps showing log expression of the DEGs across disease stages for CN10. (Likelihood ratio test, ** Benjamini Hochberg FDR < 0.01; * Benjamini Hochberg FDR < 0.05). (**F-H**) RNAscope images of *Chl1* mRNA expression in CN3/4 (n for WT= 376; n for SOD1G93A= 319), CN10 (n for WT= 377; n for SOD1G93A= 206), and CN12 (n for WT= 530; n for SOD1G93A= 508), at P112 with quantification of signal intensity (Scale bars: 30 μm, permutation test ns: P> 0.05, *: P ≤ 0.05, **: P ≤ 0.01, ***: P ≤ 0.001; ****: P ≤ 0.0001).

### Motor neuron subpopulations show unique spatial and temporal gene regulation in response to mutant SOD1 expression and resilient neurons appear rather unphased by disease

To retrieve mechanistic insight into the selective vulnerability and resilience of particular motor neuron groups in ALS, we analyzed differential gene expression induced by mutant *SOD1G93A* overexpression (Supplemental Table S1-S9). Strikingly, the resilient groups, CN3/4 and CN10 regulate very few genes with disease compared to the vulnerable CN12 and spinal motor neurons (Fig. 2A).

Analysis at the presymptomatic P56 stage, demonstrates that each motor neuron population has a largely unique response to mSOD1 (Fig. 2B, Supplemental Fig. S3A. At the onset of symptoms P112 stage, most DEGs were still unique to the particular motor neuron group. Resilient CN3/4 and CN10 neurons only share *mSOD1* as a DEG and thus clearly do not show a common induced resilience signature to ALS. However, the vulnerable CN12 and spinal motor neurons show a larger overlap in DEGs at the onset of symptoms than at the presymptomatic stage, with an overlap of 34 genes in total (Fig. 2C, Supplemental Fig. S3A).

### Upregulation of a few genes with known neuroprotective properties is sufficient for ocular motor neurons to cope with disease

To understand the spatially restricted temporal gene expression changes in disease we analyzed the DEGs within each motor nucleus across the P56 and P112 time points. Starting with the resilient motor neurons, we identified 13 DEGs unique to CN3/4 at the presymptomatic stage which recovered to baseline at the symptom onset time e.g. *Gabrq* (gamma-aminobutyric acid type A receptor subunit theta) (Fig. 2D, Supplemental Fig. S3B). Notably, *Gabrq* levels were pronouncedly higher in CN3/4 and CN10 than in vulnerable motor neuron groups, which may regulate their excitability (Supplemental Fig. S3B). In CN3/4, seven genes were dysregulated across disease stages, including; *Ucn* (urocortin) (Pedersen et al. 2002), *Cck* (cholecystokinin) (Wozniak et al. 2021) and the ECM glycoprotein *Postn* (periostin) (Shimamura et al. 2012), which all have known neuroprotective properties. *Postn* was not only upregulated in CN3/4 motor neurons with disease (Fig. 2D), but also shows several fold higher expression level in CN3/4 motor neurons than all other groups at baseline in control mice (Supplemental Fig. S3C). *Cartpt* (CART prepropeptide), *Dlk1* (delta like non-canonical notch ligand 1), *Scg2* (secretogranin II) (Fig. 2D) and *SOD1G93A* (not shown) are the remaining shared CN3/4 DEGs. Twenty genes were uniquely regulated in CN3/4 motor neurons at the symptom onset stage. Notably, *Chl1* which was reduced in CN3/4 showed the opposite regulation in CN12 and spinal motor neurons (Fig. 2D). For CN10, 43 DEGs were identified at P56 and 15 DEGs at P112, with no overlap with CN3/4 in terms of gene regulation (Fig. 2E).

We used RNAscope fluorescence *in situ* hybridization to validate gene expression changes. A probe detecting *Vacht* (vesicular acetylcholine transporter) was used to identify motor neurons. The signal was systematically quantified in each motor neuron identified from the *Vacht* channel using an automated CellProfiler pipeline (Stirling et al. 2021, v.4.2.5)(Supplemental Fig. S4A).

The downregulation of *Chl1* at P112 in CN3/4 motor neurons identified through RNA sequencing was confirmed (Fig. 2F) (one-tail mean permutation test, *P*= < 0.05), as was the lack of regulation in CN10 motor neuron (Fig. 2G) and the upregulation in CN12 motor neurons at the same time point (Fig. 2H, one-tail mean permutation test, *P*=<0.001, the number of motor neurons quantified, Supplemental Fig. S4B).

To identify if any cellular pathways were dysregulated with disease in the resilient motor neuron populations, we conducted a comprehensive analysis employing three distinct generations of pathway analysis methods: Overlap-Based Analysis (OVA) utilizing the EASE method, Per Gene Analysis (PGA) employing fGSEA, and Network Analysis (NA) with the ANUBIX method. Each method offers unique insights into the respective biology, collectively contributing to a global understanding. No pathways were significantly changed in CN3/4 motor neurons with disease, while in CN10 motor neurons four pathways were dysregulated at P112 (Supplemental Fig. S3D, Supplemental Table S10-S18). In conclusion, based on the DEGs identified in resilient neuron groups it is evident that there is no common induced resilience signature across neuron types. Some of the DEGs identified, including *Ucn*, *Cck* and *Postn*, point to neuroprotective pathways that may explain some of the tolerance towards mutant *SOD1G93A*. Nonetheless, with so few transcriptional changes identified in resilient neurons exposed to mutant SOD1 it appears conceivable that part of their resilience is encoded in their baseline gene expression.

### Identification of an innate resilience code of CN3/4 neurons and its partial induction in vulnerable motor neurons with SOD1-ALS

The combination of few gene regulation changes occurring in resilient neurons, and no pathways identified in CN3/4 motor neurons in response to *mSOD1*, prompted us to investigate the general basal gene expression signature unique to resilient neurons in control mice. We reasoned that this may explain why resilient neurons do not necessarily need to modulate their transcriptomes to a large extent in response to toxic gene expression. We found that the vast majority of transcripts were shared across resilient CN3/4 and vulnerable spinal motor neurons, as expected. CN3/4 motor neurons had 1218 uniquely detected transcripts shared across the two time points and an additional 782 transcripts unique to P56 and 733 genes unique to P112, which we reasoned could underlie their relative resilience to ALS (Fig. 3A). The transcripts uniquely enriched in either motor neuron population were visualized using Volcano plots (Fig. 3B,C, Supplemental Table S19-S20). Spinal motor neurons showed an enrichment of e.g. *Hox8-10* family members, *Mmp9*, *Dcn, Trhr and Uts2*, while CN3/4 motor neurons were enriched for e.g. *Phox2b*, *Tbx20*, *Eya1*, *Gabra1*, *Gabrb2, Glra2, Mt-Tq, Dlk1, Chrm1* and *Ucn* (Fig. 3B,C; Supplemental Fig. S3E,F and S5). This cell type specific marker enrichment is in line with previously published studies (Hedlund et al. 2010; Kaplan et al. 2014; Nichterwitz et al. 2020). We subsequently analyzed GO terms enriched in either motor neuron group at P112, and found that 18 GO terms were enriched in CN3/4 neurons (NES > 0), while 260 were enriched in spinal motor neurons (NES <0, Fig. 3D, Supplemental Table. S21-S22. To further define CN3/4 enriched gene expression valid across species, we compared with several published transcriptomics data sets, Allodi et al. 2019 (GSE93939; LCM-seq of human control CN3/4 and spinal motor neurons), Brockington et al. 2013, (GSE40438; microarray of human control CN3/4 and spinal motor neurons), Kaplan et al. 2014 (GSE52118; microarray of P9 control mice CN3/4 and spinal motor neurons) and Nizzardo et al. 2020 (GSE115130; LCM-seq of human ALS CN3/4 and spinal motor neurons). The initial analysis showed that all the data sets shared only 17 DEGs (Supplemental Fig. S6A,B). The RNA sequencing data sets revealed a substantially larger number of DEGs between CN3/4 and spinal motor neurons compared to the microarray analyses (Supplemental Fig. S6A), and we thus focused our further comparisons on the RNA sequencing data sets. Our cross-comparison with the Allodi et al. 2019 RNA sequencing data set on human *post mortem* control CN3/4 motor neurons and spinal motor neurons identified 178 CN3/4-enriched DEGs and 151 spinal motor neuron-enriched DEGs across species (Fig. 3E). Pathway analysis, revealed five shared CN3/4-enriched GO terms, including ‘Synaptic signaling’ and ‘Positive regulation of Cation Channel activity’ (Fig. 3F) and six enriched in spinal motor neurons across species, including e.g. ‘Anterior Posterior Pattern specification’ and ‘Skeletal System development’ (Fig. 3E). To try to dissect the neuroprotective program innate to CN3/4 motor neurons, we analyzed which CN3/4-enriched transcripts in baseline were induced also in vulnerable motor neurons with ALS. The reasoning being that vulnerable neurons induce transcript for protection when in distress. This analysis highlighted that vulnerable spinal motor neurons induce a set of CN3/4-baseline transcripts in response to the SOD1 mutation, including *En1*, *Gal*, *Pvalb, Gap43*, *Glra2, Cd63, Rgs10, Mt-Tq* and *Rmst* (grey circles, Fig. 3G, green box). We also found that several of these transcripts which can be considered neuroprotective, were enriched in CN3/4 motor neurons in human control *post mortem* tissues, including *EN1*, *CD63, PVALB,* and *MT-TQ* (black circles in Fig. 3G green box).

**Fig. 3.**
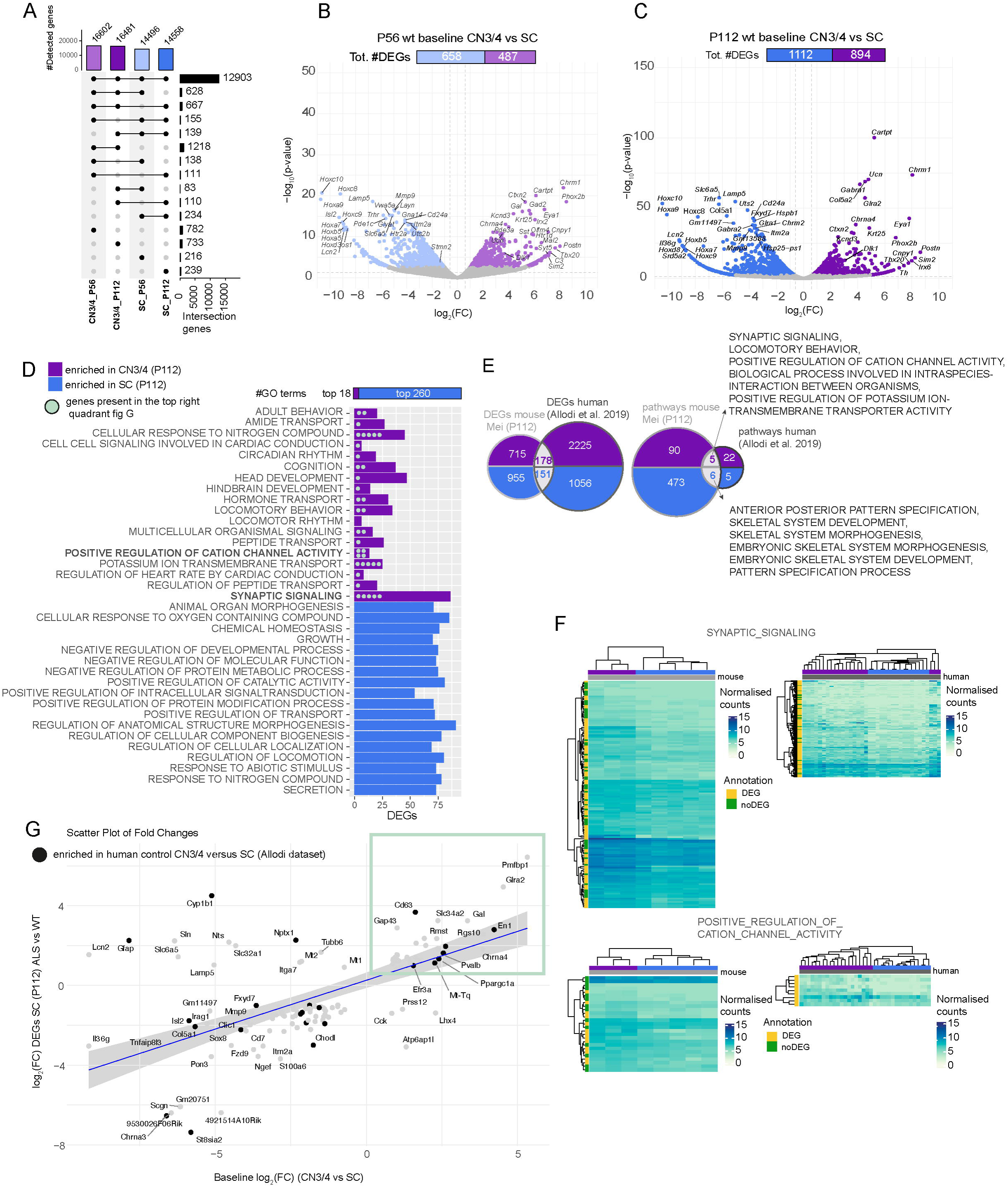
Baseline gene expression in CN3/4 motor neurons may hold the key to their resilience. (**A**) Upset plot showing the number of detected genes in CN3/4 and SC in WT mice at P56 and P112. (**B**) Volcano plot of DEGs at P56 comparing CN3/4 (light purple) versus SC (light blue), highlighting genes enriched in each population (Likelihood ratio test, Benjamini Hochberg adjusted p value < 0.05). (**C**) **Volcano plot of DEGs at P112** comparing CN3/4 (purple) vs. SC (blue), showing an increased number of differentially expressed genes at the symptom onset stage. (Likelihood ratio test, Benjamin Hochberg adjusted p value < 0.05). (**D**) Gene Ontology (GO) term enrichment analysis performed using fGSEA showing enriched GO terms in WT CN3/4 or WT spinal (SC) motor neurons at P112. (**E**) Comparison of differentially expressed genes (DEGs) and enriched pathways between mouse (Mei_P112 dataset) and human (Allodi et al., 2019). (**F**) Heatmaps of selected pathways and genes enriched in control CN3/4 motor neurons mice and in humans. (**G**) Scatter plot comparing ALS-induced expression changes in SC (P112) with baseline gene expression differences between CN3/4 and SC, identifying genes that may contribute to CN3/4 resistance. Each dot corresponds to a gene, genes labeled in bold represent those also identified in the human baseline dataset (Allodi et al., 2019). The blue line represents the linear regression of fold changes, with the shaded region showing the 95% confidence interval. Highlighted genes in the green box exhibit higher expression in CN34 baseline and induced in our SCP112, making them potential candidates for further investigation.

In conclusion, our complementary analysis of the baseline gene expression in oculomotor versus spinal motor neurons highlights their fundamental differences on top of which gene expression shifts occur with disease, mainly in vulnerable neurons. Our analysis clearly points out that basal cell type-specific gene expression needs to be considered in addition to differential gene expression with disease. In particular, our analysis highlights a set of genes and pathways that are enriched in CN3/4 motor neurons in the baseline state (without disease) and that are in part induced in vulnerable spinal motor neuron in response to disease. These are presumed to contribute to a protective response, which is maintained at a high level in resilient neurons without an apparent need for further regulation there, and insufficiently induced in vulnerable motor neurons in an attempt to keep them at bay in ALS.

### Vulnerable motor neurons show unique regulation of injury response genes indicative of cellular stress, tissue remodeling and motor neuron subtype switching

We next set out to fully dissect how relatively vulnerable motor neuron subpopulations respond to disease and the extent of temporal and spatial overlap in gene regulation. Our initial analysis clearly demonstrates that gene dysregulation in vulnerable neurons is tightly regulated both in time and space. Thus, CN12 and spinal motor neurons show mainly transcriptional responses that are unique to the cell type and disease state (Fig. 4A). Thus, to comprehensively understand how disease progresses it is necessary to conduct longitudinal analysis as it is difficult to predict the steps that will follow. Furthermore, one cell type does not necessarily inform about another, pressing the point that it is pivotal to study disease across cell types. Nonetheless, we reasoned that some of the shared responses across time may give deep insight into continuous disease predictors and drivers.

**Fig. 4.**
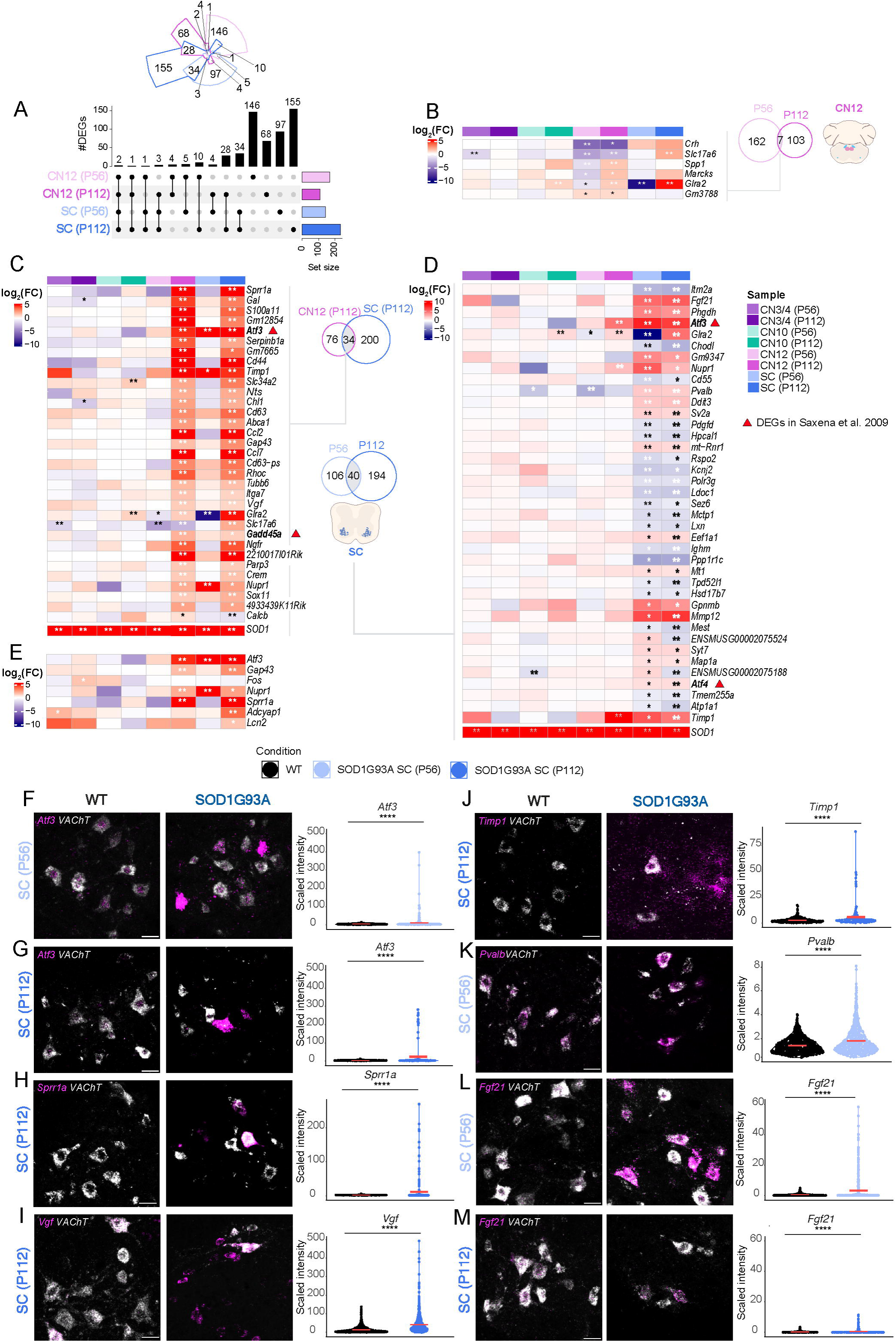
Vulnerable motor neurons show unique regulation of injury response genes. (**A**) Upset plot showing DEGs in vulnerable populations (CN12 and SC) at P56 and P112 (Likelihood ratio test, Benjamini Hochberg adjusted p value < 0.05). (**B**) Heatmap showing logFC expression of shared DEGs between ages in CN12. (**C**) Heatmap showing logFC expression of common DEGs between CN12 and SC at P112. (**D**) Heatmap showing logFC expression of common DEGs between ages in SC motor neurons. (**E**) Heatmap showing logFC expression of DEGs related to inflammation. (**B-E**) Bolded genes represent DEGs also dysregulated in Saxena et al. 2009. Likelihood ratio test, ** Benjamini Hochberg FDR < 0.01; * Benjamini Hochberg FDR < 0.05. Representative RNAscope images with quantification of signal intensity of *Atf3* in SC at P56 (n for WT= 407; n for SOD1G93A= 281) (**F**) and P112 (n for WT= 340; n for SOD1G93A= 136) (**G**), *Sprr1a* in SC at P112 (n for WT= 321; n for SOD1G93A= 218) (**H**), *Vgf* in SC at P112 (n for WT= 1031; n for SOD1G93A= 704) (**I**) *Timp1* in SC at P112 (n for WT= 340; n for SOD1G93A= 136) (**J**), *Pvalb* in SC at P56 (n for WT= 606; n for SOD1G93A= 523) (**K**), Representative RNAscope image of *Fgf21* in SC at P56 (n for WT= 592; n for SOD1G93A= 508) (**L**) and P112 (n for WT= 434; n for SOD1G93A= 258) (**M**). (**F**-**M**)(Scale bars: 30 μm, permutation test ns: P> 0.05, *: P ≤ 0.05, **: P ≤ 0.01, ***: P ≤ 0.001; ****: P ≤ 0.0001).

For CN12, only 7 of 169 DEGs from P56 were DEGs also at P112, however only half of these genes were regulated in the same direction with time (Fig. 4A,B). But of these, *Glra2* and particularly *Slc17a6*, which has neuroprotective capacities (Steinkellner et al. 2018) were also induced in spinal motor neurons at the symptom onset stage, and may be part of a protective response to damage. Interestingly, P112 CN12 motor neurons showed a higher overlap in gene dysregulation with same age spinal motor neurons than with the P56 CN12 motor neurons, indicative of a shared response to disease across vulnerable populations at the later stage. To investigate this general vulnerability code, we analyzed the 34 DEGs shared at P112 (Fig. 4C). We identified a number of genes uniquely upregulated with disease across vulnerable motor neurons, but not in resilient neurons, including e.g. *Atf3*, *Cd44*, *Gadd45a*, *Ngfr*, *Ccl2*, *Ccl7*, *Gal*, *Timp1*, *Nupr1*, *Serpinb1a*, *Ch1*, *Vgf*, *Sprr1a* and *Gap43* (Fig. 4C,D, Supplemental Fig. S4G (*Vgf*)). For spinal motor neurons, 40 DEGs were shared across the time points, while 106 DEGs were unique to the presymptomatic P56 time point and 194 DEGs to the symptom onset P112 stage, including *Nupr1*, *Atf4*, *Ddit3*, *En1*, *Gap43*, *Chl1*, *Cd44*, *Rhoc, Tubb6*, *Timp1*, *Mmp9*, *Glra2*, *Gabrg2*, *Penk*, *Chrna4* and *Dlg4* (Supplemental Table S6-7, Supplemental Fig. S4H (*Penk*)). Some of the 40 DEGs shared across time points were uniquely upregulated in spinal motor neurons (and not in CN12 motor neurons), including *Fgf21*, *Mt1, Sv2a, Chodl, Gpnmb, Mmp12, Syt7, Map1a, Atf4, Atp1a1*, *Ddit3* and *Pvalb* (Fig. 4D). *Chodl*, which is a known marker of FF motor neurons (Enjin et al. 2010) was consistently downregulated with disease in spinal motor neurons, while *Sv2a,* a marker of slow motor neurons, was upregulated (Fig. 4D), suggesting ongoing compensatory processes across motor neuron subtypes early on in disease. On the other hand, transcripts belonging to ongoing programmed neuronal death were only regulated in the later stages in spinal motor neurons (only at P112, but not at P56) and not seen in CN12 including *Adcyap1* and *Lcn2* (Fig. 4D, E), suggesting that at P56 spinal motor neurons are not at an advanced stage of neuronal damage and neither are CN12 motor neurons at P112.

To validate genes enriched in the vulnerable CN12 and/or spinal motor neurons in ALS we used RNAscope fluorescence *in situ* hybridization. A *Vacht* probe was again used to identify motor neurons. The upregulation of *Atf3* in vulnerable motor neurons (Fig. 4C) was confirmed by RNAscope at both the presymptomatic stage (Fig. 4F; one-tail mean permutation test, *P*= < 0.00001 at P56) and at the symptom onset stage (Fig. 4G; one-tail mean permutation test, *P*= < 0.00001 at P112). *Atf3* was also upregulated in vulnerable CN12 motor neurons (Supplemental Fig. S7B, D, one-tail mean permutation test, *P*= < 0.0172 at P56). While the symptom onset time point was not significant, some neurons were completely filled with *Atf3* mRNA, which was never seen in control (Supplemental Fig. S7D). *Atf3* was not upregulated in CN3/4 motor neurons at either time point (Supplemental Fig. S7A, C), in concordance with the RNA sequencing data. Consistent with axonal sprouting occurring alongside axonal degeneration, *Gap43* expression was increased in both vulnerable populations at the symptom onset stage (Fig. 4C-E). *Sprr1a* was also found upregulated in both vulnerable populations, concordant with a response to axon damage of these neurons in ALS (Fig. 4C-E) and RNAscope on spinal cord sections confirmed the upregulation with disease at the symptom onset stage (Fig. 4H, one-tail mean permutation test, *P=*< 0.0001). In CN3/4 motor neurons *Sprr1a* was undetectable, as expected (Supplemental Fig. S7E), while it was strongly induced in some CN12 motor neurons at P112, confirming the RNA sequencing result (Supplemental Fig. S7F, one-tail mean permutation test, *P=*< 0.0001).

*Vgf* was induced in both vulnerable populations at P112 and this upregulation was confirmed by RNAscope in spinal motor neurons (Fig. 4I, one-tail mean permutation test, *P=*< 0.00001). We also confirmed the upregulation of *Timp1* in spinal (Fig. 4J, one-tail mean permutation test, *P=*< 0.0001), and CN12 (Supplemental Fig. S7H, one-tail mean permutation test, *P=*<0.0001) motor neurons at P112, while it remained unchanged in CN3/4 motor neurons (Supplemental Fig. S7G) consistent with the LCM-seq data, and indicative of ongoing tissue remodeling.

*Pvalb*, which was upregulated in spinal motor neurons and downregulated in CN12 motor neurons with disease (Fig. 4D) was also confirmed by RNAscope at the presymptomatic stage (Fig. 4K, one-tail mean permutation test, *P=*< 0.00001; Supplemental Fig. S7J, one-tail mean permutation test, P<0.0001). *Pvalb* was also slightly downregulated in CN3/4 motor neurons with disease as analyzed by RNA scope (Supplemental Fig. S7I, one-tail mean permutation test, P<0.0001), following the pattern of regulation seen in the RNA sequencing data (Fig. 4D). *Fgf21* was confirmed upregulated in spinal motor neurons at P56 (Fig. 4L, one-tail mean permutation test, *P=*< 0.00001) and P112 (Fig. 4M, one-tail mean permutation test, *P*=7 e-7). *Fgf21* was also slightly upregulated in both CN3/4 and CN12 motor neurons according to the RNA scope analysis (Supplemental Fig. S7K, L, one-tail mean permutation test, *P*= 0.0187 for CN3/4 and P<0.0001 for CN12). In the RNA seq data this difference was not significant, but showed a trend towards an increase (Fig 4D). Thus, all the *in situ* data from six different probes were concordant with our RNA sequencing analysis, showing regulation in SOD1G93A motor neurons.

### Vulnerable motor neuron subpopulations degenerate in a similar fashion but at distinct temporal paces

To elucidate the pathways activated in vulnerable motor neurons, we conducted a comprehensive enrichment analysis (EA) employing methods from three different categories: OVerlap Analysis (OVA), Per-Gene score Analysis (PGA) and Network Enrichment Analysis (NEA). We selected EASE (Hosack et al. 2003; OVA), fGSEA (Korotkevich et al. 2021; PGA) and ANUBIX (Castresana-Aguirre et al. 2020; NEA), being representative of their category. Pathways showing enrichment were identified using an FDR cutoff of 0.1. The pathways demonstrating the most robust enrichment, and thus consensus across the three methods, included e.g. ‘regulation of neuronal death‘, ‘inflammatory response‘, ‘regulation of ERK cascade‘, ‘MAPK cascade‘, ‘regulation of cell adhesion‘, ‘cell migration’ and ‘synaptic signaling’ (Fig. 5A, B). Further analysis demonstrated that six of the major affected pathways were driven by 11 genes (*Atf3, Cd44, Gadd45a, Ngfr, Ccl2, Ccl7, Gal, Timp1, Nupr1, Serpinb1a* and *Chl1*) (Fig. 5C). To investigate commonality in gene network activation across vulnerable neurons, we analyzed the DEGs shared in spinal and CN12 motor neurons at P112 using Funcoup 5 (Alexeyenko and Sonnhammer, 2009; https://funcoup.org/search/). This analysis demonstrated connectivity between *Gadd45*, *Ngfr*, *Atf3* and *Cd44* in the MAPK cascade and integration with *Chl1*, in the ‘Cell Neuron Death pathway’, *Nupr1* in the ‘Cell Death pathway*‘* and *Serpinb1a* in the ‘Cell inflammation pathway’ (Fig. 5D). Moreover, using the DEGs related to common detrimental pathways in CN12 and SC at P112, we saw a clear temporal divergence along PC1 in PCA plot (Fig. 5E).

**Fig. 5.**
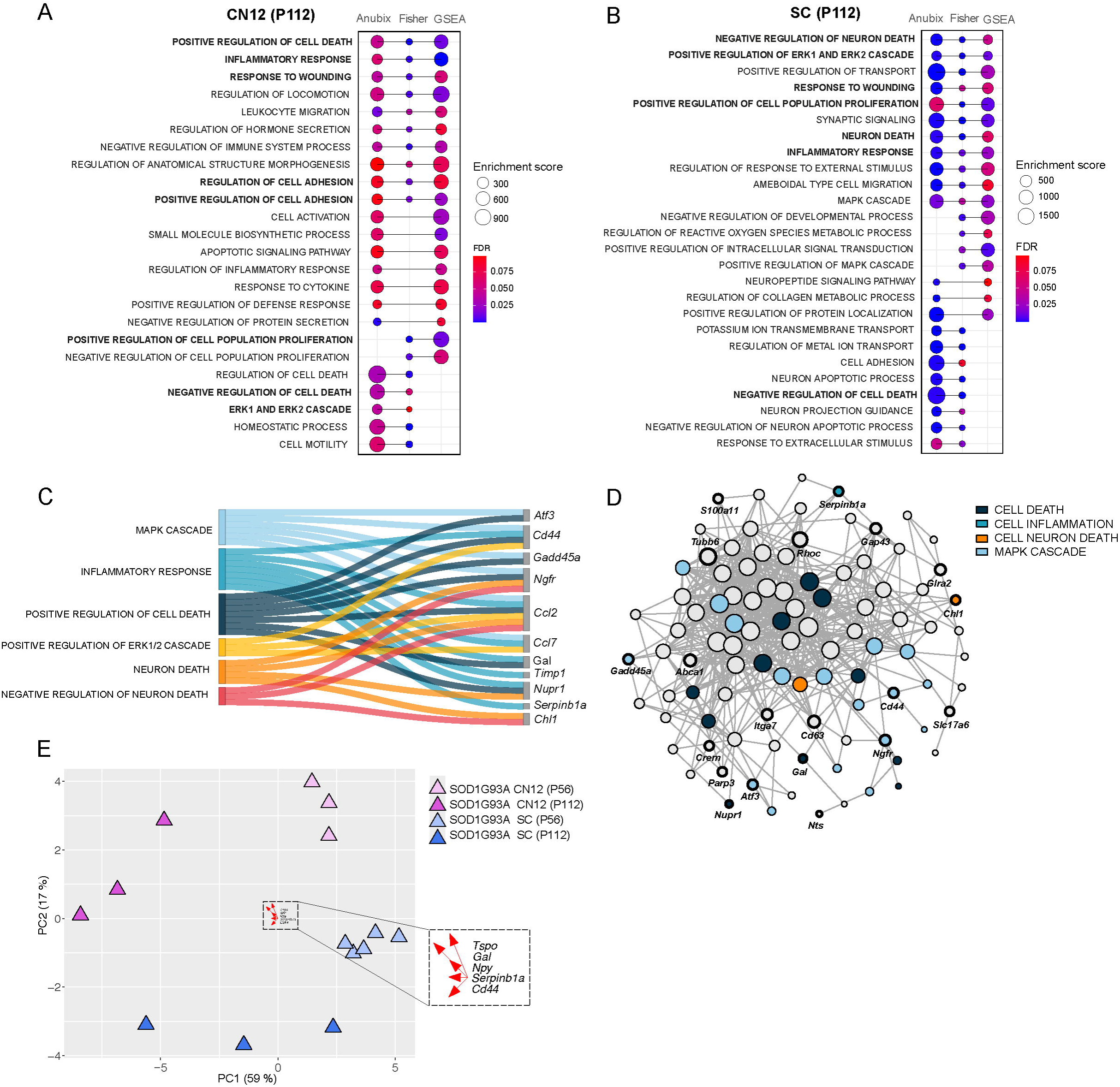
Pathway analysis shows a common detrimental response across vulnerable populations. (**A,B**) GO enrichment dot plots showing significantly enriched pathways that were enriched in at least two of the pathway analysis methods (Fisher test, fGSEA, Anubix) for CN12 and SC at P112. Enrichment scores correspond to the amount of functional genes that the method shows being related for the enrichment term. FDR threshold < 0.1. (**C**) Sankey plot showing functional categorization of stress-related genes in CN12 and SC at P112. (**D**) Functional network analysis on genes shared between CN12 and SC at P112 using Funcoup 5 with all evidence types. Evidence types are the signals that support or contradict the presence of functional coupling. In Funcoup 5 the evidences included are: Domain interactions, Genetic interaction profile similarity, Gene Regulation, mRNA co-Expression, MicroRNA Regulation, Protein co-Expression, Phylogenetic Profile Similarity, Physical Interaction, SubCellular Localization, Transcription Factor Binding profile. (E) PCA plot illustrates the clustering pattern of samples based on DEGs belonging to common detrimental pathways (‘Positive regulation of cell death’, ‘Inflammatory response’, ‘Response to wounding’, ‘Negative regulation of cell death’, ‘ERK1 and ERK2 cascade’) for CN12 and SC cell types at P112. Top 5 loading genes for PC1 are highlighted in the plot.

Our results so far reveal that there is a stress gene signature which is shared among vulnerable neurons, as resilient neurons do not show this upregulation. This signature is to a large extent shared across different vulnerable neurons, but the timing is distinct in their response to disease, with CN12 motor neurons showing a later onset and response to disease than spinal motor neurons. This result is similar to what *Saxena et al.* demonstrated for S versus FF motor neurons, where the less vulnerable S motor neurons showed a similar response to the more vulnerable FF, only later in the disease process (Saxena et al.2009). Several of the markers identified there, including *Atf3*, *Atf4* and *Gadd45a* overlap with our screen on vulnerable motor neurons (Fig. 4C, D). The lower overlap of DEGs in CN12 over time compared to spinal motor neurons likely reflects that these neurons are not as advanced in the disease process. As a result, general detrimental and regenerative processes are not activated at the presymptomatic stage, but only at the symptom onset stage. In contrast, these processes are activated across all disease stages in spinal motor neurons. This analysis solidifies our finding that CN12 motor neurons are not as far along in the disease process as spinal motor neurons in the symptomatic phase but progressing on a similar path.

### Machine learning and meta-analysis across SOD1 mutations and models identify strong disease predictors

Next, we set out to reveal which of the 129 upregulated DEGs identified in spinal SOD1G93A motor neurons at P112 (Fig. 2A) would be the strongest disease predictors. Towards this purpose we used a single cell RNA sequencing data set, *Namboori et al. 2021,* on induced pluripotent stem cell (iPSC)-derived neurons harboring a *SOD1E100G* mutation (or corrected control iPSCs), at a time when the motor neurons were starting to degenerate in cell culture (Fig. 6A). From this RNA sequencing data set we selected motor neurons (N=115) only, based on the co-expression of *SLC18A3* and *ISL1*. We used a random forest classifier (Breiman. 2001) as our machine learning approach to evaluate if our DEGs could classify these motor neurons into ALS or control (Fig. 6B). Our classification model achieved an average sensitivity of 76.0% (*P*=0.13), specificity of 73.3% (*P*=0.05), PPV of 50.3% (*P*=0.08), NPV of 90.9% (*P*=0.12), AUC of 84% (*P*=0.16) and accuracy of 74.0% (*P*=0.03). These classification performance measures were obtained using a rigorous nested Leave-One-Out (LOO) cross-validation approach, which ensures that the evaluation of performance is based on left-out samples rather than the training data. These results demonstrate the effectiveness of our model in distinguishing between different genotypes using our upregulated DEGs in SCP112. The genes with the strongest importance for the classification were *VGF*, *PENK*, *NTS (neurotensin)* and *INA (internexin neuronal intermediate filament protein alpha)* (Fig. 6C). Other genes contributing to the classification, *SV2A* and *GAP43,* indicate motor neuron subtype switching and/or enrichment with disease and regeneration, respectively.

**Fig. 6.**
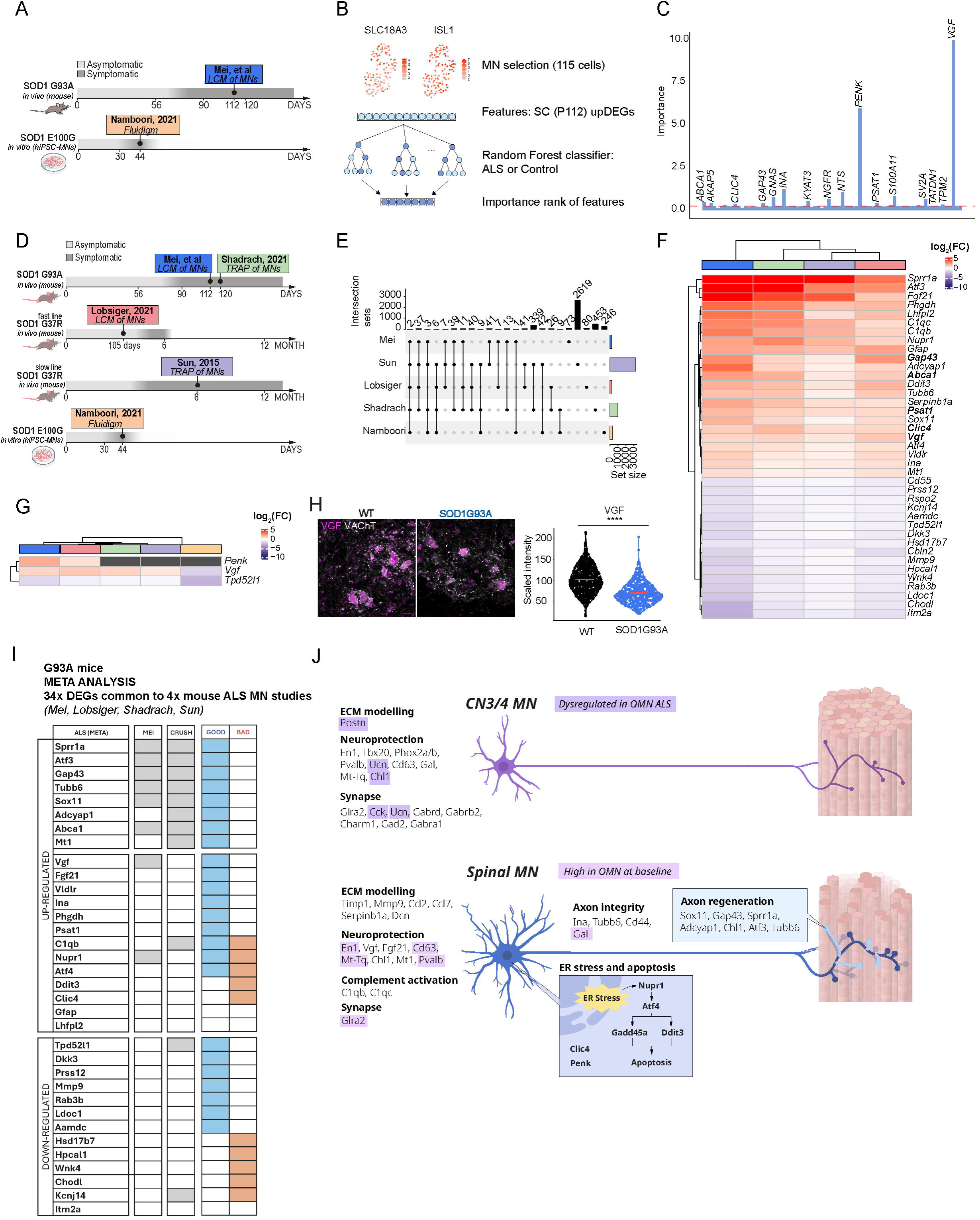
Dataset comparison and feature selection show a common importance for Vgf in ALS. (**A**) Overview of the comparison between two datasets: Namboori et al. 2021 and SC (P112) dataset (**B**) Schematic of the random forest-based machine learning approach for identifying key ALS-relevant genes using upregulated DEGs in SC (P112) (n cells selected in Namboori et al.: 115). (**C**) Feature importance ranking from random forest classifier, highlighting Vgf as the most predictive marker. (**D**) Overview of cross-dataset DEG comparison, including Shadrach et al. 2021, Lobsiger et al. 2007, Sun et al. 2015, Namboori et al. 2021. (**E**) Upset plot showing DEGs shared across ALS datasets (Likelihood ratio test, Benjamini Hochberg adjusted p value < 0.05). (F) Heatmap of common DEGs identified across all mouse studies between SC at P112, Shadrach et al. 2021, Sun et al. 2015, Lobsiger et al. 2007. Bolded gene names are also predictive in the radom forest disease classifier. (**G**) Heatmap showing common DEGs between all datasets. (**H**) Representative immunofluorescence image of SC at P112 and showing the expression of VGF protein in VAChT positive cells, Scale bar: 30 μm. (**I**) Heatmap displays 34 ALS-induced DEGs common to four mutant SOD1 motor neuron (MN) studies (Mei, Lobsiger, Shadrach, and Sun). Genes are categorized as upregulated or downregulated in ALS motor neurons. The presence of each gene in different datasets is indicated in columns: MEI (disease-induced DEGs shared between CN12 and SC motor neurons at P112 from Mei et al.), CRUSH (sciatic nerve crush model from Shadrach et al., 2021) and if the regulation is considered GOOD or BAD. (**J**) Summary of transcriptional differences in baseline gene expression between resilient and vulnerable neurons as well as their differential responses to disease, organized into functional categories, reveal key pathways including ER stress and apoptosis, axon integrity and regeneration, neuroprotection, synapse, complement activation and ECM remodeling.

We next evaluated if these disease-predicting DEGs were commonly regulated in spinal motor neurons in response to mutant *SOD1* across data sets. We reasoned that shared regulation across mouse cohorts, mutations, time points and even species would reveal important disease regulators. Such a meta-analysis across models and SOD1 mutations has not been done previously and is clearly challenging, as there is a temporal regulation in gene expression with disease, as seen in our SOD1G93A data. Nonetheless, we did also identify overlap across time points particularly in the vulnerable neuron groups, speaking to the feasibility of such an analysis. We thus analyzed three additional published data sets that isolated RNA from spinal motor neurons from mice overexpressing human mutant *SOD1* as well as the Namboori et al. sequencing data on human stem cell-derived neurons harboring a point mutation in *SOD1*; *i*) *Lobsiger et al. 2007* isolated motor neurons from lumbar spinal cord from onset of symptoms (15 weeks of age) in a fast progressing SOD1G37R mouse (reaches end-stage at 6 months) using LCM and Affymetrix Arrays; *ii*) *Sun et al. 2015* used TRAP to isolate mRNAs on ribosomes in motor neurons at symptom onset (8 months of age) in a slow progressing SOD1G37R mouse (reaches end-stage at 12 months) and *iii*) *Shadrach et al. 2021* used TRAP to isolate RNA on ribosomes in motor neurons in symptomatic SOD1G93A (4 months of age) (Fig. 6D). The four mouse studies, including the present one, shared 39 DEGs (Fig. 6E,F). Among these were *Vgf*, *Gap43*, *Ina, Psat1, Clic4* and *Abca1* (Fig. 6F), that all aided in the classification of human motor neurons into ALS or control (Fig. 6C). *Penk*, which was a strong disease predictor in the human data set (Fig. 6C) was identified in our data and that of *Lobsiger et al. 2007* (Fig. 6G). Only two DEGs were found across all five studies (mouse and human) and those were *Vgf and Tpd52l1* (Fig. 6G). While only upregulated DEGs were used to classify the human data set (Fig. 6C) *Tpd52l1* was consistently downregulated across all data sets, and thus may also predict disease (Fig 6F). *Vgf* was also upregulated in CN12 with disease, but remained unaffected in resilient neuron groups (Fig. 3B). Furthermore, it should be noted that while *Vgf* was upregulated in all four mouse studies, it was down-regulated in *Namboori et al. 2021*. (Fig. 6G). A previous study has shown that the VGF protein level in cerebrospinal fluid (CSF) was decreased in ALS patients by approximately 40% and VGF immunoreactivity in *post mortem* spinal motor neurons was also decreased (Zhao et al. 2008). As we saw the opposite pattern on the mRNA level across all mouse studies, we decided to analyze the VGF protein level using immunofluorescence staining of SOD1G93A spinal cords and wild-type littermates (N=5/genotype). As for the RNA scope analysis, the signal was quantified in each motor neuron identified from the Vacht channel using an automated CellProfiler pipeline (Supplemental Fig. S4A). It demonstrated that VGF protein was indeed decreased in ALS motor neurons with disease (Welch’s two-sample t-test P= <2.2e-16, Fig. 6H), consistent with previous findings in patients (Zhao et al. 2008). Thus, the mRNA increase of *Vgf* may be a compensatory event due to the loss of VGF protein. To get a comprehensive understanding of all common DEGs across the mouse studies and their implications for disease we compared them to the published results on RNA sequencing of motor neurons after nerve crush (Shadrach et al. 2021), that would delineate regenerative versus detrimental responses (Fig. 6I). We clustered gene regulation as either “GOOD” or “BAD” responses for ALS. Thus, if a neuroprotective/regenerative gene was down-regulated, it was classified as a “BAD” response. This analysis clearly indicates that ALS motor neurons do not simply degenerate, but that there are protective compensatory processes activated along with the deleterious. Overall “GOOD” responses included neuroprotective genes that were up-regulated: *Sprr1a, Atf3 and Gap43* mediating pro-regenerative responses; *Phgdh/Psat1* (L-serine pathway), and protective responses against stress that were up-regulated e.g. *Adcyap1* and *Mt1*. It also included downregulation of detrimental genes including e.g. *Mmp9* (vulnerability marker) and *Ldoc1* (pro-apoptosis). In contrast, overall detrimental (BAD) responses included up-regulation of pro-apoptosis genes *Clic4* and *Nupr1*, and downregulation of neuroprotective genes *Cbln2, Cd55 (DAF)* and *Rspo2* (Fig. 6I). Finally, to further interpret and summarize our findings, both on differences found in baseline gene expression between resilient and vulnerable neurons as well as their responses to disease, we investigated functional categories curated across multiple studies, which revealed key pathways linked to ER stress and apoptosis, axon integrity and regeneration, neuroprotection, synapse, complement activation and ECM remodeling (Fig. 6J). In conclusion, we have identified a resilience code in ALS, as well as strong ALS disease markers and predictors based on DEGs identified in our study and across multiple published data sets, which predict disease across SOD1 mutations and species.

## Discussion

We conducted RNA sequencing analysis of motor neurons displaying differential vulnerability to degeneration in ALS to unveil mechanisms of protection and degeneration at distinct disease stages. Our analysis did not reveal any major differences mutant *SOD1* mRNA levels across neuron types or in other disease-associated gene expression that could explain their differences in vulnerability. Thus, the cause for differences in susceptibility to ALS lies elsewhere and analysis of responses induced by mutant *SOD1* may give mechanistic insight into both why certain neurons succumb to disease and why others persist.

Analysis of differential gene expression consistently showed that resilient motor neurons display a very mild transcriptomic response to disease while vulnerable motor neurons showed large transcriptional dysregulation already early in disease. This is different from the gene dysregulation we previously noted in spinal muscular atrophy (SMA), where resilient CN3/4 motor neurons elicited a large early gene response compared to vulnerable spinal and facial (CN7) motor neurons which only responded to the loss of the *Smn1* gene (survival motor neuron 1) later in disease. In the SMA mouse model we saw an induction of a large set of intuitively protective genes in CN3/4 motor neurons in combination with a disease module that was shared across vulnerable and resilient neurons (Nichterwitz et al. 2020). This indicates that CN3/4 motor neurons are not as severely affected by misfolded mutant SOD1 as they are to loss of SMN and consequent disruption of RNA splicing. It is also consistent with CN3/4 motor neurons showing less aggregation of misfolded SOD1 compared to vulnerable spinal motor neurons or CN12 motor neurons, as demonstrated in the SOD1G93A mouse (An et al. 2019) and in SOD1G85R mice (Thomas et al. 2018.)

Even though SOD-ALS resilient neurons, both CN3/4 and CN10, regulated very few genes and thus did not seem in need to modulate gene expression in a major way to survive, the gene expression changes that did occur may hold a partial key to their resilience. Seven genes were dysregulated across disease stages in CN3/4 motor neurons, including *Ucn* (urocortin), *Postn* (periostin), *Cartpt* (CART prepropeptide), *Cck* (cholecystokinin), *Dlk1* (delta like non-canonical notch ligand 1), *Scg2* (secretogranin II) and *mSod1*. Of these DEGs *Ucn* has been shown to protect hippocampal neurons from oxidative stress and excitotoxic glutamate insult at picomolar levels (Pedersen et al. 2002). *Cck* has been shown to protect Purkinje cells from toxicity in ataxias (SCA mice) (Wozniak et al. 2021). *Postn* (periostin) levels were not only upregulated in CN3/4 motor neurons with disease, but were also several fold higher in CN3/4 motor neurons than all other groups at baseline in control mice. POSTN is an ECM glycoprotein that can stimulate neurite outgrowth (Shih et al. 2014) and be neuroprotective as shown in cortical neurons (Shimamura et al. 2012). To better comprehend why CN3/4 motor neurons do not need to regulate a large number of genes to cope with disease, we conducted a complementary analysis of the baseline expression differences between CN3/4 and spinal cord motor neurons in control across time. This analysis highlights the fundamental differences between somatic motor neuron types.

Here it became evident that while CN3/4 motor neurons expressed a somewhat larger number of genes, albeit many of those at low levels, the number of GO terms uniquely enriched in each nucleus was overwhelmingly larger in spinal motor neurons with 260 enriched GO terms compared to only 18 in CN3/4 motor neurons. Thus, oculomotor neurons may require fewer processes for their function than spinal motor neurons do, maybe due to their lower metabolic demand and output onto many fewer muscle fibers.

Our cross-comparison with neurons isolated from human *post mortem* tissues identified a number of pathways unique to each motor neuron group which was conserved across species, including the CN3/4 enriched GO terms of ‘Synaptic signaling’ and ‘Positive regulation of Cation Channel activity’. To further visualize the innate neuroprotective program of CN3/4 motor neurons we analyzed which CN3/4-enriched transcript were induced in vulnerable motor neurons with ALS, with the hypothesis that vulnerable neurons induce transcript for protection when in distress. Our analysis highlights that several CN3/4-baseline transcripts, including *En1*, *Pvalb*, *Gap43*, *Glra2, Gal, Cd63, Rgs10* and *Rmst* are induced in vulnerable motor neurons with SOD1-ALS. Of these transcripts, *EN1*, *CD63, PVALB* and *MT-TQ* and were also relatively enriched in CN3/4 motor neurons compared to spinal motor neurons in *post mortem* tissues from control patients. This indicates that baseline gene expression differences that may explain CN3/4 motor neuron resilience is shared across species. It also highlights En1 as a potential CN3/4-resilience factor, which may benefit from further potentiation in vulnerable neurons to improve their resilience to ALS. In fact, En1 is known to act as a neuroprotective factor to motor neurons. In the spinal cord, En1 is normally not produced by motor neurons, but rather by V1 spinal interneurons, that synapse on alpha-motor neurons, deliver this homeodomain protein paracrinely. If this delivery is blocked motor neurons start to degenerate (Leboeuf. et al. 2023).

Our finding that En1 mRNA is induced within vulnerable motor neurons in ALS clearly demonstrate an attempt at neuroprotection, similar to what CN3/4 motor neurons display already at baseline without being challenged, and maintain throughout disease.

Our analysis also pinpoints that to understand vulnerability and resilience we need to examine baseline gene expression difference in connection with analyzing disease-induced dysregulation.

In addition to DEG analysis, we also made a first attempt to examine polyadenylation in our data. To this end we found that both CN3/4 and spinal motor neurons show large changes in polyadenylation with disease, a response that was partially overlapping across populations and time, but also showed unique regulation related to the particular motor neuron group in question and the disease stage (Supplemental Fig. S8A-H). This opens up a new interesting avenue for future research using complementary approaches, to investigate the fate of particular modified transcripts to elucidate if this leads to changes in subcellular localization of transcripts, their splicing and/or half-life of the resulting proteins.

Several of the genes regulated in vulnerable spinal motor neurons were previously shown to be upregulated in spinal motor neurons in mutant SOD1 mice (Lobsiger et al. 2007). Our study reveals that this gene signature is unique to vulnerable neurons, as resilient neurons do not show this upregulation and that it is shared across SOD1 mutations, and is clearly part of a stress response to disease. Vulnerable spinal motor neurons showed an upregulation of a gene set known to be important for nerve regeneration, including *Gap43*, *Cd44*, *Chl1, Atf3, Sprr1a* and *Adcyap1*. This highlights, that vulnerable motor neurons during ALS are not just dying, but that they actively try to overcome the ongoing degeneration, by inducing genes that stimulate regeneration. It has also been shown that certain spinal motor neurons are able to sprout and compensate for the muscle denervation of neighboring motor neurons, leading to a temporary stabilization of NMJs (Frey et al. 2000; Fischer et al. 2004; Schaefer et al. 2005). This results in a temporary stabilization of NMJ loss versus gain (Comley et al. 2016) which is thought to slow the decline in motor dysfunction. Reduction of Cd44 has been shown to reduce axon initiation of retinal ganglion cells (Ries et al. 2007) and thus an increase as seen in the SOD1G93A motor neurons is expected to promote axon growth. The upregulation of *Atf3* in spinal motor neurons across time points and at the late time point in CN12 is also likely protective as it has been shown that overexpression of *Atf3* promotes motor neuron sprouting and survival as well as retained innervation of muscle in ALS mice (Seijffers et al. 2014). *Sprr1a* is known to be an axon regeneration-induced gene (Starkey et al. 2009), suggesting a role in protective responses. Similarly, the upregulation of *Pvalb* is anticipated to be protective, as it can safeguard neurons from excitotoxicity (Van Den Bosch et al. 2002) as well as Gal, which also has neuroprotective properties, as shown for hippocampal neurons (Elliott-Hunt et al. 2004).

The large majority of motor neuron somas are still present at the symptom onset stage we examined (Chiot et al. 2021). Thus, the downregulation of *Chodl*, an FF motor neuron marker in spinal motor neurons and upregulation of the S motor neuron marker Sv2a, could be indicative of adaptation across these two populations. It may also be an indication of subtype switching among motor neurons. FF motor neurons are highly vulnerable to ER stress and start to show patterns of denervation early in disease (Saxena et al. 2009) while S motor neurons compensate by sprouting in the mSOD1 mouse (Pun et al. 2006). Innervation of FF muscle groups by S motor neurons will induce muscle subtype switching towards an S type. As muscle also talks back to motor neurons and affect their identity (Correia et al. 2021) this may also impact FF motor neurons as these try to reconnect with their now modified muscle target, but this remains to be further investigated. Nonetheless, some of the unique regulation in spinal motor neurons at the symptom onset may also represent a slight difference in proportion of motor neuron subtypes incorporated (during the LCM isolation) rather than an induction of gene regulation. Future single cell or nuclei RNA sequencing studies may be able to further resolve this issue if a temporal analysis is conducted.

The upregulation of *Syt7* in only spinal motor neurons is indicative of their need to reorganize presynaptic function, while the upregulation of *Lcn2* on the other hand appears to be a way to negatively control regeneration (Gu et al. 2024), perhaps a way to block motor neurons that are already handling too much stress- and DNA damage responses from regenerating, but this remains to be further investigated. Hypoglossal (CN12) motor neurons also showed an induction of regenerative genes including *Gap43*, *Cd44* and *Chl1*. Some transcripts belonging to a disease response were only regulated in the later stages in spinal motor neurons and not seen in CN12, including *Adcyap1*, which is also linked to nerve regeneration (Baskozos et al. 2020). This may both be a reflection of that CN12 motor neurons are not as far along into the disease process as spinal motor neurons and that they have cell-type specificity in their response to disease. To fully resolve this question, neurons isolated from later stage diseased animals would need to be studied.

The consequences of the upregulation of *Ccl2* and *Ccl7* in vulnerable neurons, a response that is shared across ALS and nerve injury, as shown by Shadrach and colleagues (Shadrach et al. 2021), is not completely clear. Literature indicates that it could lead to enhanced neuronal excitability (Zhou et al. 2011), and glial activation in the CNS, (Joly-Amado et al. 2020), which may be detrimental.

*Fgf21* was robustly and specifically upregulated in spinal motor neurons across disease stages. FGF21 is involved in regulating carbohydrate and lipid metabolism and maintaining energy homeostasis, and it can protect cells from apoptosis. FGF21 is induced by ER stress, mitochondrial dysfunction and starvation. We and others have recently shown that ALS motor neurons display mitochondrial dysfunction early on (Hor et al. 2021; Mehta et al. 2021; Schweingruber et al. 2023), and thus *Fgf21* upregulation may be a consequence thereof. The upregulation of *Fgf21* in vulnerable spinal motor neurons may also be indicative of the increased energy demand on these cells during disease. There is a clear correlation between increased serum levels of FGF21 and metabolic disease conditions such as diabetes, mitochondrial diseases, obesity and aging, all of which have muscle loss as a common factor (reviewed in Tezze et al. 2019).

The upregulation of (nuclear protein 1) in CN12 and spinal motor neurons is indicative of the stress these cells are experiencing, as this transcriptional regulator is involved in ER stress, oxidative stress response, DNA repair, autophagy, apoptosis and chromatin remodeling (*reviewed in* Liu et al. 2022). In spinal motor neurons it was regulated already at the presymptomatic stage while in CN12 neurons only at the symptom onset stage, demonstrating that these cell types follow a similar path at distinct paces, where spinal motor neurons have taken the lead to destruction. Furthermore, *Ddit3* (DNA-damage inducible transcripts 3) (CHOP) is an ER-stress apoptotic mediator which was upregulated in spinal motor neurons alone, which is another clear indication that these cells are further along a degenerative pathway than CN12 motor neurons. *Ddit3* mRNA was previously shown to be upregulated in spinal motor neurons in 90-120 day-old SOD1G93A mice (Perrin. et al. 2005), and at 15 weeks in the SOD1G37R mouse (Lobsiger et al. 2007) as well as the protein level in end-stage sporadic ALS patient motor neurons (Ito et al. 2009). The earlier detection in our data, already at P56, may be a reflection of the sensitivity of Smart-seq2 and is in concordance with the very early ER stress response of fast fatigable (FF) motor neurons, seen prior to any visible denervation (Saxena et al. 2009). The upregulation of *Mt1* in spinal motor neurons across time points indicate a response to block apoptosis (Zhang et al. 2013), and *Gpnmb* appears to be part of an inductive protective response, as this molecule can be protective in ALS and is upregulated in sera from sporadic ALS patients (Tanaka et al. 2012).

Extracellular matrix remodelling is regulated by the activity of MMPs (matrix metalloproteinases) which are tightly regulated by tissue inhibitors (Timps). The upregulation of *Timp1* (metallopeptidase inhibitor 1) across time points in spinal motor neurons and in the symptomatic stage in CN12 motor neurons, indicate again that these two vulnerable motor neuron groups follow similar paths of response and destruction albeit on a different time scale. *Mmp12* was also upregulated in spinal motor neurons indicating ongoing ECM remodeling.

In general, we note that many gene regulations in vulnerable neurons indicate compensatory events to handle the toxicity of mutant SOD1. The majority of these appear to be beneficial, such as nerve regeneration programs, but clearly insufficient over time. We also note that different vulnerable neuron populations share some responses to ALS but that their timing is distinct, likely due to differences in their temporal involvement in the disease.

To further understand the strength in the identified vulnerability signature in both defining and predicting disease across data sets, we compared our RNA sequencing data to that of five other published studies. We found that *Vgf* and *Tpd52l1* were identified as DEGs across all data sets independent of disease status, presymptomatic, or symptom onset stage, and across SOD1G37R, SOD1G93A and SOD1E100G mutations. A 4.8 kDa VGF peptide was previously described as a potential biomarker of ALS as the levels were decreased in CSF of ALS patients and could distinguish them from controls (Pasinetti et al. 2006). VGF is involved in energy expenditure so decreased levels have been hypothesized to contribute to the hypermetabolic state seen in ALS. A follow-up study confirmed VGF as a biomarker for ALS using ELISA which indicated that VGF CSF levels may be correlated with muscle weakness in ALS patients (Zhao et al. 2008). While *Vgf* mRNA was upregulated in our data and all other mouse transcriptome studies analyzed, we found VGF protein to be decreased in ALS motor neurons with disease in ALS mice, consistent with previous studies. The inverse correlation between RNA and protein levels indicates that there is a compensatory mechanism at play to increase VGF either by increased transcription, or by changes in RNA stability.

In conclusion we demonstrate that resilient motor neurons only regulate a few genes in response to mutant *SOD1* likely as their baseline gene expression renders them resilient against this specific insult. The few genes that were upregulated in CN3/4 have known protective properties, including *Ucn*, *Cck* and *Postn* and may confer resilience. We also demonstrate that CN3/4 has high baseline activity of several neuroprotective genes, including *En1*, *Gal*, *Cd63* and *Pvalb* that are maintained in these neurons in ALS and specifically upregulated in vulnerable neurons in response to disease.

One key distinction between ALS-vulnerable spinal motor neurons and CN3/4 neurons lies in their ability to maintain inhibitory synaptic transmission, a crucial factor in neuronal stability and protection against excitotoxicity. In spinal motor neurons, Glra2 (glycine receptor) is lowly expressed compared to CN3/4 motor neurons, but increases in ALS at symptom onset, possibly as a delayed compensatory mechanism to restore inhibition. Despite this, the concurrent downregulation of Gabrg2 (GABA-A receptor subunit) suggests a continued weakening of inhibitory GABAergic input in these neurons. Additionally, CN3/4 motor neurons exhibit a high level of neuroprotective receptors such as Gad2, Gabrd, Gabrb2, Gabra1, which stabilize neuronal firing rates and prevent hyperexcitability. These differences in synaptic regulation, alongside the presence of neuroprotective and ECM-stabilizing factors like Postn suggest that CN3/4s employ a distinct resilience mechanism that enables their survival in ALS, contrasting with the progressive degeneration seen in spinal motor neurons.

We also reveal that different vulnerable motor neuron populations share pathway activation, which indicate that cell death occurs through similar mechanisms across vulnerable motor neurons, but temporally separated. The DEG and pathway analyses clearly demonstrate that vulnerable motor neurons activate a majority of beneficial and neuroprotective gene programs including for nerve regeneration reflecting their effort to reconnect. ALS-vulnerable spinal motor neurons exhibit increased ER stress and apoptotic signaling, with markers such as *Nupr1*, *Atf4*, and *Ddit3* promoting cell death. Additionally, C1qb/C1qc upregulation suggests that these neurons are actively marked for clearance, further driving neurodegeneration. As ALS-vulnerable motor neurons show *Timp1* upregulation and lower *Mmp9* levels we speculate that this could lead to increased ECM stiffness, due to a decreased level of ECM degradation, inflammation as well as impaired axonal plasticity.

These differences in synaptic regulation, alongside the presence of neuroprotective and ECM-stabilizing factors like Postn suggest that OMNs employ a distinct resilience mechanism that enables their survival in ALS, contrasting with the progressive degeneration seen in spinal motor neurons. Machine learning and meta-analysis across mutant SOD1 data sets and disease time points reveal a shared transcriptional vulnerability disease code and identify *VGF*, *PENK, INA*, *GAP4*3 and *TPD52l1* as strong disease-predictors across SOD1 mutations and species. These genes thus have potential as future biomarkers of disease and may aid in diagnosis and prognostics. In conclusion, our study reveals motor neuron population-specific basal gene expression and temporal disease-induced regulation that together provide a basis to explain ALS selective vulnerability and resilience. Our findings also provide further support for that ALS vulnerable motor neurons do not simply undergo degeneration, but that compensatory and neuroprotective mechanisms are at play. We reveal a number of resilience mechanisms that may provide novel therapeutic targets aimed to enhance synaptic stability and neuroprotection in vulnerable neurons.

## Methods

### Ethics statement and animal model

All procedures involving animals were approved by the Swedish ethics council, a local animal ethics committee (Paris CE5, France) and carried out according to the Code of Ethics of the World Medical Association (Declaration of Helsinki). Animals were housed with a 12-hour dark/light cycle under standard conditions and had access to food and water *ad libitum*. Adult SOD1G93A mice on a C57Bl/6J background (B6.Cg-Tg(SOD1-G93A)1Gur/J, The Jackson Laboratories strain # 004435, males) were used as a model of ALS and non-transgenic littermates served as a control. All animals were anesthetized with a lethal dose of avertin (2,2,2-Tribromoethanol in 2-Methylbutanol, Sigma-Aldrich) prior to either decapitation or intracardial perfusion with phosphate buffered saline (PBS) followed by 4% paraformaldehyde (PFA) in PBS. Animals used for RNA sequencing originated from a cohort of animals in Sweden, all animals used for RNAscope derived from a cohort located in France.

### Tissue processing and laser capture microdissection for transcriptomics

For the dissection and processing of tissues for LCM-seq, all equipment, surfaces, and tools were carefully cleaned with RNaseZap (Ambion/Life Technologies) and wiped with distilled water or 70% ethanol. Brain and spinal cord tissues were dissected from 56 and 112-day old males and immediately snap-frozen in 2-Methylbutane (Sigma-Aldrich) on dry ice, followed by storage at -80°C until further processing. After equilibration of the tissue to approximately - 20°C, 12 µm coronal sections were prepared on a cryostat and placed onto PEN membrane glass slides (Zeiss). Spinal cords were embedded in OCT (Dako) prior to sectioning. Slides were stored at -80°C until further processing. For laser capture microdissection (LCM), cells were visualized with a quick histological staining (Histogene, Arcturus/Life Technologies) followed by dehydration in a series of ethanol solutions (75%, 95%, and 99.7%, Solveco). LCM was performed on a Leica LMD 7000 system at 40x magnification and cutting outlines were drawn in close proximity to individual cells to minimize contamination by surrounding tissue. Per sample, approximately 100-200 cells with an area of >200 μm^2^ (150 μm^2^ for vagus motor neurons) and a visible nucleus with nucleolus were collected into the dry cap of a PCR tube (Biozym Scientific). After the addition of 5 µl lysis buffer (0.2% Triton X-100, with 2 U/μl recombinant RNase inhibitor, Clontech) to the cap, samples were mixed by pipetting up and down, spun down in a tabletop centrifuge and snap frozen on dry ice. Samples were stored at -80°C until library preparation.

### Tissue processing for immunohistochemistry and RNAscope

Male and female mice were transcardially perfused with PBS followed by 4% PFA in PBS at P56 or P112. Brains and the lumbar region of the spinal cord was dissected, post-fixed for 3 h in 4% PFA at 4°C, and cryoprotected in PBS 30% sucrose for 48 h at 4°C. The lumbar region of the spinal cords and the brains were sectioned at 30 μm thick using Thermo Scientific Sliding Microtome Microm HM 430. For the RNAscope experiments we used three control and three mutant SOD1 mice for spinal cord at P112 and brains regions at P56 and P112. For the spinal cord at P56 we used four mice for control and ALS.

### cDNA and sequencing library preparation

For RNA-seq experiments, all reagents were of molecular biology/PCR grade if available. Only nuclease-free water (H_2_O, LifeTechnologies) and tubes that were certified nuclease-free were used. All workbenches and equipment were cleaned with RNaseZAP (Ambion/Life Technologies) and additionally with DNAoff (Takara) for library preparations.

For library preparation, a modified version of the Smart-seq2 protocol (Picelli et al. 2014b, 2013) was used, which is described in detail in (Nichterwitz et al. 2018, 2016). All heat incubation steps and PCR cycles were carried out in a BioRad T100 Thermal Cycler. After reverse transcription, cDNA was amplified for 18 cycles, followed by purification with magnetic beads (GE Healthcare). cDNA concentration and library quality were assessed using an Agilent 2100 Bioanalyzer (High Sensitivity DNA kit). Three samples with poor cDNA library quality were excluded from analysis (low cDNA concentration with no apparent peaks). One ng of cDNA (as determined with the Bioanalyzer, 100-9000 bp range) was used as input for the tagmentation reaction, which was carried out with 0.4 -1 μl of in house Tn5 (Picelli et al. 2014a). Ligation of sequencing indices (Nextera XT Sequencing Index Kit, Illumina) and 10 cycles of enrichment PCR were performed with Kapa HiFi polymerase. Final sequencing libraries were purified with magnetic beads and the concentration of each sample was determined on a Qubit fluorometer (ThermoFisher) using the dsDNA high sensitivity kit (LifeTechnologies). Samples were pooled and sequenced on the Illumina Hiseq 2500 seq platform.

### RNA-seq analysis

The RNA-seq reads were mapped simultaneously to the mm39 mouse genome assembly and the genomic sequence of human SOD1 from the hg38 assembly using STAR (version 2.7.0e) (Dobin et al. 2013). The majority of reads across samples were uniquely mapped to the mouse genome, with a minority being multi mapped or non-mapped (Supplemental Fig. S2A). We used only uniquely mapped reads for further analyses. Expression levels were determined using the rpkmforgenes.py software (http://sandberg.cmb.ki.se/rnaseq) with the Ensembl gene annotation. Samples included in the analysis had more than 15,000 detected genes (16408 ± 75 genes; *mean ± SEM*). Genes with at least one count in at least five samples were retained in the dataset. Before Differential Expression Analysis (DEA), samples were normalized using the calcNormFactors function from the edgeR package. This function employs the trimmed mean of M-values (TMM) method by default, which normalizes raw count data by accounting for differences in library sizes and other systematic biases. Following normalization, the dispersion was estimated and the model was fitted using a quasi-likelihood (QL) approach, which provides robust error rate control for differential expression analysis. Differential expression analysis was performed using the R-package EdgeR (v3.36.0 Robinson et. al. 2010). A quasi-likelihood negative binomial generalized log-linear model was fitted to the data before empirical Bayes moderated t-statistics were calculated and multiple testing correction (Benjamin-Hochberg) was performed. The threshold false discovery rate (FDR) < 0.05 was used to determine if a transcript was differentially expressed. No minimum fold change was applied to identify differentially expressed genes. For Supplemental Fig. S2D, an ANOVA was implemented with one factor (Cell type_ Age) which was significant. We then ran post-hoc analyses to assess pairwise differences and t-test results with Bonferroni correction are reported.

### Enrichment analysis

The Enrichment analysis methods used in this study belong to three different categories: OVerlap Analysis (OVA), Per-Gene score Analysis (PGA) and Network Enrichment Analysis (NEA) methods. As methods, we selected EASE (OVA), fGSEA (Korotkevich et al. 2021, PGA) and ANUBIX (Castresana-Aguirre M et al. 2020, NEA). NEA methods stand out from other approaches due to their consideration of the interconnections among DEGs within the context of functional gene sets in a network framework (Buzzao et al. 2024). All methods were run in an R (v4.1.1) version-controlled conda environment using the original packages, EASE and ANUBIX were run as in a recent benchmarking study for gene set EA methods (Buzzao et al. 2024). EASE and ANUBIX were performed on differentially expressed genes with an FDR below 0.1. FunCoup (Alexeyenko and Sonnhammer, 2009; https://funcoup.org/search/) was used as a network with a default confidence link score of 0.8 and 5 max neighbors in order to obtain the network of links that exist between vulnerable genes within detrimental pathways.

Gene Set Variation Analysis (Hänzelmann S. et al. 2013, GSVA, v.1.50.1) was performed to assess pathway enrichment. GSVA was used on log-transformed counts per million (CPM) values calculated from the expression data. A GSVA parameter object was then created with the ‘gsvaParam’ function, using log-transformed CPM values, list of GO-BP gene sets (MSigDB), kcdf=Gaussian and maxDiff=False. The ‘gsva’ function was subsequently applied to compute enrichment scores for each sample.

### Random forest classifier

To classify ALS and control samples, we implemented a Random Forest (RF) model using the ranger package in R. The input gene expression dataset from Namboori et al. (2021) was preprocessed using DESeq2 normalization (v.1.34.0, Love et al., 2014), followed by z-score transformation using the scale() function, ensuring that each gene’s expression values were centered around the mean and scaled to unit standard deviation. To address class imbalance between ALS (n = 30) and WT (n = 85) samples, we applied downsampling using the downSample() function from the ROSE package, ensuring an equal distribution of classes during training.

We employed nested cross-validation (CV) to optimize the RF model and prevent overfitting. The outer CV loop (5-fold) repeatedly split the dataset into training and test sets, while the inner CV loop (5-fold) was used for hyperparameter tuning via the train() function from the caret package. The RF model was trained using the following hyperparameter grid: mtry {5, 10, 15, 20} (number of randomly selected features at each split), splitrule “gini” (Gini impurity for node splitting), and min.node.size {2, 7, 12} (minimum number of samples per terminal node). The best-performing model was selected based on its predictive performance within the inner CV loop.

The outer CV loop evaluated the generalization performance of the model using the left-out test sets, assessing accuracy, sensitivity, specificity, positive predictive value (PPV), negative predictive value (NPV), and area under the ROC curve (AUC). Additionally, we computed permutation-based feature importance to assess the contribution of individual genes to disease classification. To establish statistical significance, we conducted permutation testing (n = 100 iterations) by randomly shuffling disease labels and re-evaluating model performance. The resulting p-values for accuracy, sensitivity, specificity, PPV, NPV, and AUC were computed by comparing the observed performance to the permutation-derived null distribution.

### APA quantification

The MAAPER software (v. 1.1.1) was used to map sequencing reads to known polyadenylation (PA) sites, as specified in the PolyA DB v3 database for the mm39 genome (Li et al. 2021; Wang et al. 2018). To detect genes exhibiting significant alterations in the length of their 31-most exon, we applied the REDu metric provided by MAAPER. REDu quantifies the relative expression levels of the two most differentially expressed isoforms in the 31-most exon. Positive REDu values signify transcript lengthening, while negative values indicate shortening events. To control for multiple testing, we applied the Benjamini-Hochberg (BH) correction to the REDu p-values, obtaining FDR-adjusted p-values (FDR_pval). Genes with FDR_pval < 0.05 were considered significantly altered in APA.

### Use of published datasets

To evaluate the purity of our LCM collected samples, we compared our data to a previously published dataset obtained from GEO, accession number GSE52564 (Zhang et al. 2014), which was processed as described for our samples. For analysis of oculomotor versus spinal motor neuron enriched transcripts across species we compared with our published RNA sequencing data set from Allodi et al (GSE93939) and Nizzardo et al (GSE115130), with Brockington et al. 2013 (GSE40438) and Kaplan et al. 2014 (GSE52118). For the comparison of our data with other spinal motor neuron microarray and RNA-seq data, the previously published data were preprocessed as described below. The microarray data (Lobsiger et al. 2007) were analyzed after rma() normalization and quality control for both moe430A and B arrays. Shadrach et al. 2021 and Sun et al. 2015 were mapped to the mm39 assembly, using STAR (version 2.7.0e). Genes with a minimum count of 4 and 3, respectively, within all samples were retained. Namboori et al 2021, was mapped to the hg38 assembly and processed retaining all cells (n=115 cells) with co-expression of *Slc18a3* and *Isl1* of at least 5 counts and nUMI >= 1000, nGene >= 300 and mitoRatio<= 0.5). Differential analysis was carried out using Wald test in DESeq2 and parameters set as follows: sfType = “poscounts”, minReplicatesForReplace = Inf, useT = TRUE and minmu = 1e-06. We chose to use DESeq2 for the differential expression analysis for the Namboori dataset, to ensure comparability and consistency, as this methodology was used in the original paper, with the reported results (DESeq(dds, test = “Wald”, sfType = “poscounts”, minReplicatesForReplace = Inf, useT = TRUE, minmu = 1e-06)). To our knowledge, there is no dedicated function specifically designed for single-cell differential expression analysis in EdgeR that provides comparable optimization possibilities to DESeq2 without resorting to pseudo-bulking.

### RNAscope fluorescent *in situ* hybridization

As previously described (Leboeuf et al. 2023), sections were selected by region of interest (lumbar region for the spinal cords and CN3/4, CN10 and CN12 for the brains). They were washed in PBS, incubated with RNAscope hydrogen peroxide solution from Advanced Cell Diagnostics (ACD) for 10 min at room temperature (RT), rinsed in PBS 0.1% Tween 20 at RT, collected on Super Frost plus microscope slides (Thermo Scientific), dried at RT for 11h, rapidly immersed in ultrapure water, dried again at RT for 11h, heated for 11h at 60°C, and dried at RT overnight. The next day, sections were immersed in ultrapure water, rapidly dehydrated with 100% ethanol, incubated at 100°C for 15 min in RNAscope 1X Target Retrieval Reagent (ACD), washed in ultrapure water, and dehydrated with 100% ethanol for 3 min at RT. Protease treatment was carried out using RNAscope Protease Plus solution (ACD) for 30 min at 40°C in a HybEZ oven (ACD). Sections were then washed in PBS before *in situ* hybridization using the RNAscope Multiplex Fluorescent V2 Assay (ACD). Probes were hybridized for 2 h at 40°C in a HybEZ oven (ACD), followed by incubation with signal amplification reagents according to the manufacturer’s instructions. Probes were purchased from ACD: Mm-Slc18a3-C3 (catalog #448771-C3), Mm-Atf3-C2 (catalog #426891-C2), Mm-Sprr1a-C2 (catalog #426871-C2), Mm-Timp1-C1 (catalog #316841), Mm-Fgf21-C1 (catalog #460931), Mm-Pvalb-C4 (catalog #421931-C4), Mm-Vgf-C2 (catalog #517421-C2), Mm-Chl1-C1 (catalog# 531051). The hybridized probes’ signals were visualized and captured on a Zeiss LSM800 Airyscan confocal microscope with a 20× objective.

### RNA scope Image analysis and quantification

All mouse spinal cord sections from ages P56 and P112 were imaged at 20X on a confocal microscope (Axio-Observer Z1/7). Scans were acquired at high resolution (4084 × 4084) with a z-step size of 0.58 µm. Maximum intensity images were firstly generated from merged z-focal planes using CellProfiler (Stirling et al. 2021, v.4.2.5). Intensity of RNAscope probes and immunohistochemistry images were quantified within the masked *Vacht*/ChAT+ cells using an automated CellProfiler pipeline. Each cellular specimen exhibiting an intensity of *Vacht*/ChAT+ signal higher than 0.01 was subjected to quantification. Each image, after being z-projected using Cellprofiler, was processed following the steps below: The images were imported as grayscale. The main approach started applying median filtering to the *Vacht* channel of the images using the ‘MedianFilter’ module. We calculated illumination correction with the ‘CorrectIlluminationCalculate’ module and then applied it using the CorrectIlluminationApply module for all the channels of the image (see Supplemental Figure 4A). This step allowed correction for possible illumination unbalance. We then enhanced the features signal of the *Vacht* channel to identify motor neurons better. We then used the ‘IdentifyPrimaryObjects’ module to identify the motor neurons and mask them, employing per-object thresholding and manual adjustments to the smoothing filter size and maxima suppression distance to optimize segmentation. Following this, we measured the intensity of the masked motor neurons for all the other channel probes we were interested in with the ‘MeasureObjectIntensity’ module and assessed their size and shape using the ‘MeasureObjectSizeShape’ module. We placed outlines on the images using the ‘OverlayOutlines’ module and saved the processed images with the ‘SaveImages’ module. We repeated this semi-automatic pipeline for all the images of all the conditions and ages. Statistical testing was performed utilizing one tailed mean permutation test with a permutation count of 100,000 and the nominal p-value is presented.

### Spinal cord immunohistochemistry

Sections were quickly washed in PBS and incubated in 5% donkey serum, 0.3% triton-X 100 in PBS for 1 h at RT. Sections were then incubated overnight with primary antibodies diluted in 5% donkey serum, 0.3% triton-X 100 in PBS at 4°C, washed and further incubated with secondary antibodies for 1 h at RT. Primary antibodies included rabbit anti-VGF (LSBio LS-C352987, 1:100), goat anti-VAChT (Millipore ABN100, 1:500). Controls without primary antibodies were included. Images were acquired on a Zeiss LSM800 Airyscan confocal microscope with a 20× objective.

### Data access

All LCM-seq RNA-seq data generated in this study have been deposited at the Gene Expression Omnibus (GEO) of the National Center for Biotechnology Information, with the accession number GSE244538.

## Acknowledgements

And we would also like to thank all members of the Hedlund and the Bartosovic laboratories for fruitful discussions and helpful suggestions regarding lab meetings. We would like to thank Marc Friedländer for excellent discussions on quality controls in RNA sequencing. We would like to thank Chris Molenaar, facility manager at IFSU for providing excellent technical expertise on confocal imaging. We would like to thank Anna Klemm for her great assistance in the RNAscope images analysis. We would also like to thank Dr Rita Almeida (SU) for her insightful comments and suggestions regarding statistical analysis of RNAscope data. We would like to thank Mattias Karlén for his outstanding work on creating schematics in Figures 1 and 6. We would also like to thank Professor Bin Tian and Dr. Shan Yu for their valuable assistance in providing the appropriate version of the PAS database corresponding to the mouse genome version mm39. Confocal microscopy was performed at the Imaging Facility at Stockholm University (IFSU). The computations and data handling were enabled by resources provided by the National Academic Infrastructure for Supercomputing in Sweden (NAISS) and the Swedish National Infrastructure for Computing (SNIC) at UPPMAX. This work was supported by grants from the Swedish Research Council (2016-02112 and 2020-01049) to E.H., European Union Joint Programme for Neurodegenerative Disease (JPND) (529-2014-7500) to EH, from the Department of Biochemistry and Biophysics, Stockholm University to E.H., from Åhlén’s Foundation (Åhlén-stiftelsen) to E.H., from Ulla-Carin Lindquists foundation for ALS research (Ulla-Carin Lindquists stiftelse för ALS forskning) to E.H., and Birgit Backmarks Donation to ALS research at Karolinska Institute (Birgit Backmarks Donation till ALS forskning vid Karolinska Institutet) to E.H. I.L. has been financed by a MESR (’French ministry for high education and research’) PhD fellowship. Project related funding for C.S.L. comes from the french national ALS association (‘Association pour la recherche sur la SLA - ARSLA’) and additional ALS associations (’Aide à la Recherche des Maladies du Cerveau - ARMC’, and ‘SLA Fondation Recherche - SLAFR’). C.S.L. thanks the staff of the animal housing facility UMS28/CEF (Paris, France) and of the iGenSeq platform for genotyping (ICM, Paris, France; which received funding from the program ‘Investissements d’avenir’ ANR-10-IAIHU-06).

## Author contributions

E.H. conceived the study. I.M., S.N., M.L., D.L and E.H. designed experiments. E.H., D.R. and C.L. supervised the project. I.M., S.N., M.L., J.N., I.L., and C.L., acquired data, specifically S.N., and J.N., collected tissues for LCM and isolated neurons using LCM. S.N. prepared cDNA libraries for sequencing; I.M., and J.N. mapped sequencing data. I.M. conducted all bioinformatics analysis of RNA sequencing data; C.L. and I.L prepared tissues for RNA scope; M.L. conducted RNA scope; M.L. and I.M. conducted RNA scope imaging and analysis. M.L. and I.M. conducted immunofluorescent analysis. I.M., S.N., M.L., J.N., C.L. and E.H. analyzed data; I.M., M.L., S.N., E.H. prepared all Figures for paper. E.H. wrote the initial manuscript with the help of I.M., M.L. and C.L. All authors edited and gave critical input on the manuscript.

