## Supplementary Figure legends for "Transcriptional modulation unique to vulnerable motor neurons predicts ALS across species and SOD1 mutations"

**Supplemental Figure legends**

**Supplemental Figure S1.** Laser capture microdissection (LCM) of neurons from motor nuclei and spinal cord. (**A**-**D**) Representative brightfield images showing motor neuron populations before and after LCM in the oculomotor/trochlear nucleus (CN3/4) (**A**), dorsal motor nucleus of the vagus nerve (CN10) (**B**), hypoglossal nucleus (CN12) (**C**), and lumbar spinal cord (SC) (**D**). Motor neurons were stained with cresyl violet prior to dissection to enable precise visualization. (**A**’-**D’**) Close-up images of the boxed regions in A-D highlight motor neuron morphology prior to laser capture. (**A”**-**D”**) Images after LCM showing successful motor neuron removal, with the remaining tissue demonstrating cleanly excised neuronal profiles. Scale bars: A = 200 µm (applies to A, A”, B, B”, C, C”); D = 400 µm (for D and D”); D’ = 50 µm (applies to A’, B’, C’, D’).

**Supplemental Figure S2.** Positional identity and neuronal enrichment validation in motor neuron subpopulations. (**A**) Mapping statistics of RNA-seq samples (n=54) mapped to the mouse genome (mm39) with an added human SOD1 sequence. It shows the percentage of uniquely mapped, multi-mapped, and non-mapped reads per motor neuron subtype. Only uniquely mapped reads were used in downstream analyses. (**B**) Heatmap of Hox gene expression profiles, evaluating the positional identity of motor neuron populations along the rostrocaudal axis. Hox gene expression patterns confirm expected regional identity of neurons from CN3/4, CN10, CN12, and spinal cord (SC). (**C**) Heatmap of neuronal marker gene expression, confirming an enrichment of motor neurons in the dataset. LCMseq samples were compared to a previously published RNA-seq data set of neurons (black), astrocytes (orange), oligodendrocyte precursor cells (OPCs, light turquoise), newly formed oligodendrocytes (NFO, medium turquoise) and myelinating oligodendrocytes (MO, turquoise) to evaluate the purity of our neuronal samples (GEO accession number GSE52564). (**D**) Boxplots visualizing RNA levels of the human SOD1 transgene (top) and endogenous mouse Sod1 (bottom) in transgenic animals. Data indicate robust transgene expression in ALS mice and consistent levels of endogenous Sod1 across cell types. Statistical tests: one-way ANOVA with Bonferroni’s multiple comparison test (*P ≤ 0.05; **P ≤ 0.01; ***P ≤ 0.001; ****P ≤ 0.0001). (**E**) Heatmap of Gene Ontology (GO) enrichment scores from Gene Set Variation Analysis (GSVA), assessing pathways related to ubiquitin-proteasome function, protein degradation, and autophagy-related processes in CN3/4 and spinal motor neurons of control and SOD1G93A mice. (F-G) Principal component analysis (PCA) of additional PCs (PC3-6) did not show further clustering of disease versus control.

**Supplemental Figure S3. Differential gene expression of selected genes and Gene Ontology (GO) enrichment analysis for CN10.** (**A**) Upset plot showing the overlap of differentially expressed genes (DEGs) across all motor neuron subtypes and ages. This visualization highlights shared and unique DEGs across CN3/4, CN10, CN12, and SC at P56 and P112, showing common pathways between resistant and vulnerable populations. (**B-C, E-H**) Expression patterns of selected genes in different motor neuron subtypes and ages. Heatmaps (top panels) display normalized counts of each gene across all samples, while boxplots (bottom panels) show log2-transformed RPKM expression levels, highlighting differences between genotypes ( SOD1G93A vs. WT) and cell types; (**B**) *Gabrq*, (**C**) *Postn*. (**E**) *Dcn,* (**F**) *Dlk1*, (**G**) *Vgf*, (**H**) *Penk*. (**D**) GO term dot plot representing enriched biological processes in CN10 at P112. The Gene Ontology (GO) enrichment analysis was performed using three complementary generations of enrichment methods: Fisher’s exact test, functional gene set enrichment analysis (fGSEA), and Anubix. Enrichment scores correspond to the amount of functional genes that the method shows being related for the enrichment term. FDR threshold < 0.1.

**Supplemental Figure S4. Image processing and quantification workflow for RNAscope analysis. (A)** Pre-processing and segmentation workflow for RNAscope images using CellProfiler. The RNAscope probe used for VAChT/ChAT detection was GFP-labeled. The top row displays the pre-processing steps: the original image after applying a median filter (left), the illumination correction function used to normalize the image (center), and the final corrected image (right). The bottom row shows the segmentation workflow: the input image from the first analysis cycle (left), the segmented probeGFP objects highlighted in different colours (center), and the final outlines of the detected objects overlaid on the corrected image (right). The table on the bottom right provides example statistics, including the number of detected objects (n = 9), object size metrics (10th, median, and 90th percentile diameters) and the area covered by objects (2.4%). (**B**) Table showing the number of motor neurons that were quantified for the RNAscope probes used.

**Supplemental Figure S5. Validating the expression of selected DEGs in the motoneurons**

**(A-G)** We confirmed the mRNA localization of *Stmn2, Eya1, Chrm1, Chrna4, Sst, Phox2b* and *Gal* in CN3/4 (enlarged on the right panel) of p56 mice brains. **(H-N)** We confirmed the mRNA localization of *Stmn2, Hoxc10, Mmp9, Cd44, Ccl7, Trhr* and *Pde1c* in the spinal motoneurons (enlarged on the right panel) of p56 mice. (© 2004 Allen Institute for Brain Science. Allen Mouse Brain Atlas and Allen Mouse Spinal Cord Atlas. Available at: [*http://mouse.brain-map.org/*](http://mouse.brain-map.org/) and [*http://mousespinal.brain-map.org*](http://mousespinal.brain-map.org/) )

**Supplemental Figure S6. Differentially expressed genes (DEGs) in wild-type baseline datasets between ocular motor neuron (OMN; CN3/4) and spinal motor neuron (SC).** (**A**) Upset plot of DEGs across multiple datasets comparing OMN vs. SC in SCP112 (Mei et al.), Allodi et al. (2019), Kaplan et al. (2014), and Brockington et al. (2013). Mei et al. and Allodi et al. (2019) are RNAseq dataset and Kaplan et al. (2014), and Brockington et al. (2013) microarray datasets. (**B**) Heatmap of DEGs that were shared across datasets, clustered by expression profile similarity. Genes are color-coded based on expression intensity (logFC between OMN and SC), with upregulated genes in warm tones (orange-red) and downregulated genes in cool tones (blue-purple).

**Supplemental Figure S7: Representative RNAscope images and quantifications in CN3/4 and CN12 regions at different ages.** (**A-L**) Representative RNAscope images and quantifications of transcript scaled intensity (raw intensity multiply by a scaling factor of 1000) across CN3/4 and CN12 motor neurons in WT and SOD1G93A mice. Images show fluorescent in situ hybridization signals for selected genes, with corresponding quantification of fluorescence intensity per motor neuron. (**A**) Representative RNAscope image with quantification of signal intensity of *Atf3* in CN3/4 at P56 (n for WT= 221; n for SOD1G93A= 198), (**B**) in CN12 at P56 (n for WT= 488; n for SOD1G93A= 300), (**C**) in CN3/4 at P112 (n for WT= 262; n for SOD1G93A= 192), (**D**) in CN12 at P112 (n for WT= 308; n for SOD1G93A= 289). (**E**) *Sprr1a* in CN3/4 at P112 (n for WT= 376; n for SOD1G93A= 319), (**F**) and in CN12 at P112 (n for WT= 530; n for SOD1G93A= 508). (**G**) *Timp1* in CN3/4 at P112 (n for WT= 303; n for SOD1G93A= 132), (**H**) and in CN12 at P112 (n for WT= 323; n for SOD1G93A= 297). (**I**) *Pvalb* in CN3/4 at P56 (n for WT= 278; n for SOD1G93A= 131). (**J**) in CN12 at P56 (n for WT= 579; n for SOD1G93A= 238). (**K**) *Fgf21* in CN3/4 at P56 (n for WT= 221; n for SOD1G93A= 198). (**L**) and in CN12 at P56 (n for WT= 488; n for SOD1G93A= 300). (A-L) **(Scale bars: 30 μm. Quantifications performed using a permutation test; nd: not detectable (mean raw intensity < 0.8 in both conditions), ns: P> 0.05, *: P ≤ 0.05, **: P ≤ 0.01, ***: P ≤ 0.001, **: P ≤ 0.0001).

**Supplemental Figure S8. Alternative polyadenylation (APA) analysis in CN3/4 and spinal motor neurons.**

**(A, B, D, E)** Volcano plots showing differential alternative polyadenylation (APA) site usage as assessed by the REDu metric. The x-axis represents REDu (log2 fold change), while the y-axis shows the -log10 p-value. Positive REDu values indicate transcript lengthening, and negative values indicate shortening. Genes with the most significant REDu changes are labelled. (**A**) Comparison of CN3/4 at P56. (**B**) Comparison of SC at P56. (**D**) Comparison of CN3/4 at P112. (**E**) Comparison of SC at P112.
**(C, F, G, H)** Venn diagrams illustrating the overlap of genes with significant APA changes (REDU.pval < 0.05) between different conditions. (**C**) CN3/4 at P56 vs. SC at P56. (**F**) CN3/4 at P112 vs. SC at P112. (**G**) CN3/4 at P56 vs. CN3/4 at P112. (**H**) SC at P56 vs. SC at P112.
